# Zymocin-like killer plasmids were present in the common ancestor of terrestrial fungi

**DOI:** 10.1101/2025.10.13.682091

**Authors:** Padraic G. Heneghan, Letal I. Salzberg, Kenneth H. Wolfe

## Abstract

Some budding yeasts secrete killer toxins made by linear dsDNA plasmids located in the cytosol. The best-known example is the *Kluyveromyces lactis* toxin zymocin, which is encoded by a 9-kb killer plasmid assisted by a 13-kb helper plasmid. These plasmids are distantly related to eukaryotic dsDNA viruses and have been called Virus-Like Elements (VLEs) but they do not produce virus particles. Their evolutionary origin is unclear because VLEs have been found only in budding yeasts (subphylum Saccharomycotina of phylum Ascomycota) and not in any other fungi. Here, we show that similar VLEs are present in two deeply divergent phyla of terrestrial fungi, Zoopagomycota and Mucoromycota. In Zoopagomycota, some isolates of *Coemansia* harbor more than 20 different linear dsDNA plasmids simultaneously, many of which encode their own DNA polymerases (DNAPs). We did not identify any functional killer toxins encoded by the new VLEs, but phylogenetic analysis of the chitinase genes present on some of them suggests that they are orthologs of the chitinase subunits of killer toxins encoded by Saccharomycotina VLEs. Phylogenetic analysis of DNAPs shows that the diversity of VLEs in Zoopagomycota greatly exceeds that in Saccharomycotina, and that Saccharomycotina killer and helper plasmids are related to two lineages of VLEs present in Zoopagomycota. Our results indicate that VLEs were present in the common ancestor of all terrestrial fungi about 650 Mya, and that they were already subdivided into killer and helper types by this time.

## Introduction

The majority of well-studied species in the kingdom Fungi belong to the subkingdom Dikarya, consisting of the phyla Ascomycota and Basidiomycota, but other phyla that are often collectively called the Early Diverging Fungi (EDF) contain the greatest phylogenetic diversity in the kingdom and are the descendants of its deepest speciation events (Mondo and Grigoriev 2025). Only in the past decade has genome sequencing given us a relatively clear view of the number of EDF lineages and their phylogenetic relationships (Spatafora et al. 2016; Berbee et al. 2017; James et al. 2020; Amses et al. 2022; Strassert and Monaghan 2022). It has become apparent that there many separate phylum-level lineages of fungi outside the Dikarya (Fig. 1A). Consequently, terms such as EDF or ‘basal fungi’ refer to paraphyletic groups and their use has been discouraged (James and Rokas 2025). The deepest branches of fungi are zoosporic (flagellated) fungi, which are generally motile, aquatic, species (Mondo and Grigoriev 2025). They form outgroups to the terrestrial fungi, which are non-motile. The terrestrial fungi clade (Fig. 1A) includes the phyla Mucoromycota and Zoopagomycota as well as the Dikarya, and is about 650 million years old (Chang et al. 2021). Mucoromycota and Zoopagomycota both produce zygospores and were previously considered to be a single phylum (Zygomycota), but genome sequencing revealed that they are phylogenetically separate, with Zoopagomycota lying outside a clade consisting of Mucoromycota + Dikarya (Spatafora et al. 2016; Chang et al. 2021). There are several subphyla within each of the phyla of terrestrial fungi (Fig. 1A), and some aspects of the phylogeny are still not completely resolved (James et al. 2020; Amses et al. 2022; Strassert and Monaghan 2022).

**Figure 1.**
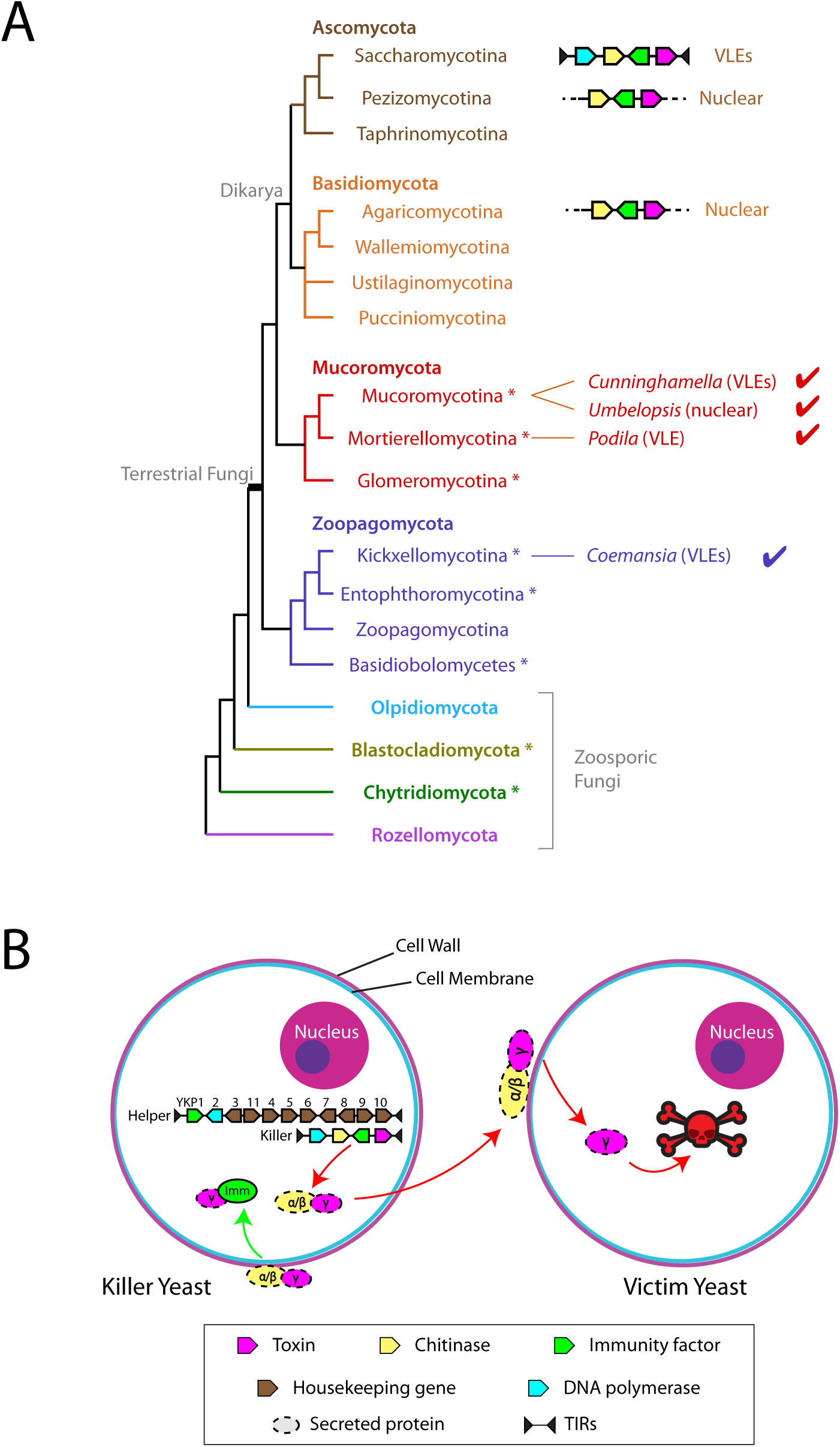
Virus-like elements in the kingdom Fungi. **(A)** Phylogenetic relationships in Kingdom Fungi. Gene structure cartoons illustrate the taxa previously known to contain linear plasmid VLEs (in Saccharomycotina) or VLE-like sequences integrated into the nuclear genome (in many Pezizomycotina species and a few Agaricomycotina species) (Heneghan et al. 2025a; Heneghan et al. 2025b). Check marks indicate the taxa in which we report VLEs in this study, in phyla Mucoromycota and Zoopagomycota. Names are in bold are phylum names. The tree topology and clade names are from Mondo and Grigoriev (2025), based on phylogenomic analyses. Lineages without reference genome sequences are not shown. Asterisks indicate taxa reported to contain NCLDV pseudogenes in their nuclear genomes (Gong et al. 2020; Myers et al. 2025). **(B)** Mechanism of action of a killer toxin such as zymocin in *Kluyveromyces lactis*. The toxin is encoded by one VLE (the killer plasmid), and a second VLE (the helper plasmid) is required for its maintenance and expression. Both of these VLEs are multi-copy linear dsDNA molecules located in the killer cell’s cytosol. The γ-subunit of the toxin kills the victim cell by cleaving a specific tRNA or rRNA. Housekeeping genes on the helper plasmid are given names beginning with *YKP*.

Fungi host almost no known viruses with double-stranded DNA (dsDNA) genomes, and none are known in subkingdom Dikarya (Gong et al. 2020; Myers et al. 2025). However, very recently, evidence of dsDNA giant viruses actively infecting several species of zoosporic fungi was reported, including the near-complete genome sequence of a virus infecting an *Allomyces* species (phylum Blastocladiomycota) (Myers et al. 2025). Giant viruses are also called nucleocytoplasmic large DNA viruses (NCLDVs) and comprise the viral phylum *Nucleocytoviricota*. They have very large genomes, for example the 1-megabase genome of the amoeba virus *Mimivirus* (Colson et al. 2017). Indirect evidence that a wide range of fungi has been infected by dsDNA giant viruses in the past has been inferred from the presence of *Nucleocytoviricota*-derived pseudogene sequences in their nuclear genomes (Gallot-Lavallee and Blanc 2017; Gong et al. 2020; Myers et al. 2025). Such pseudogenes have been detected in multiple fungal phyla and subphyla (Fig. 1A), which suggests that *Nucleocytoviricota* have an ancient and recurring association with the fungal kingdom (Gong et al. 2020).

Although they do not contain known dsDNA viruses, some budding yeasts (subphylum Saccharomycotina of phylum Ascomycota) contain linear cytosolic dsDNA molecules that are called virus-like elements (VLEs) or cytosolic plasmids (reviewed in (Satwika et al. 2012; Schaffrath et al. 2018)). The most famous of these are the killer plasmids of yeasts such as *Kluyveromyces lactis*. Killer strains of this species contain two cytosolic plasmids: a 9-kb ‘killer’ plasmid that codes for a secreted ribonuclease toxin called zymocin that can kill other yeast species, and a 13-kb ‘helper’ plasmid that contains housekeeping genes necessary for the replication and transcription of both plasmids (Fig. 1B). The toxin consists of a rapidly-evolving ribonuclease (called the toxin γ subunit, or simply γ-toxin), which is secreted and delivered to the victim cell by a chitinase protein (called the α/β subunit), where it kills the victim by cleaving tRNA or rRNA. As well as coding for the toxin subunits, killer VLEs also code for an immunity protein that is not secreted and protects the killer cell from its own toxin (Fig. 1B). The VLEs replicate in the cytosol, independently of the nuclear genome, and are slightly beneficial parasites because the toxin enables their hosts to kill competitors (Wickner and Edskes 2015). VLEs are rare, being present in only 1-2% of yeast isolates, and have not been found outside the budding yeast subphylum Saccharomycotina (Fukuhara 1995; Heneghan et al. 2025a). Some yeast isolates contain helper plasmids without having killer plasmids, and others contain ‘cryptic’ plasmids that are non-autonomous (i.e., they require a helper plasmid to be present) but do not appear to code for a zymocin-like killer toxin (Satwika et al. 2012; Polomska et al. 2021; Heneghan et al. 2025a).

The yeast cytosolic plasmids are called VLEs because some of their genes show a clear phylogenetic relationship to virus genes, even though they do not produce virus particles (Satwika et al. 2012). Some of the housekeeping genes of VLEs (i.e., two RNA polymerase subunits, a helicase, and an mRNA capping enzyme) appear to have originated from *Nucleocytoviricota* (Krupovic and Koonin 2015). However, the DNAP polymerase (DNAP) genes of VLEs are related to the DNAPs of a different group of dsDNA viruses, subphylum *Polisuviricotina* (adenovirus, virophages, polintons, and their relatives) (Koonin et al. 2024). This apparently dual origin led to the proposal that VLEs descended from a recombination event between *Nucleocytoviricota* and *Polisuviricotina* ancestors, followed by an evolutionary reduction in which the proto-VLE lost the ability to make a virus particle and adopted a non-viral mode of propagation (Krupovic and Koonin 2015; Krupovic et al. 2024).

*Saccharomycotina* VLEs replicate by using a DNAP encoded by the VLE itself (Arzumanyan et al. 2018). These polymerases are in the pPolB family (also called B-type DNAPs) which initiates replication by using a protein primer, unlike most other DNAPs which use an RNA primer. The protein primer, known as the terminal protein (TP), is covalently attached to the 5’ ends of the linear plasmid DNA and provides an anchor point for the DNAP to begin replication. The DNAP and TP are encoded by a single gene in the VLE, synthesized as a precursor DNAP-TP fusion protein that is later cleaved to yield separate DNAP and TP (Takeda et al. 1996; Krupovic et al. 2024). The same mechanism is found in *Polisuviricotina* viruses (Koonin et al. 2024). Structural analyses of the DNAPs have confirmed the relationship between the yeast VLEs and these dsDNA viruses (Krupovic et al. 2024). However, to date, VLEs have been found only in Saccharomycotina, a phylogenetic space exceptionally distant from where their viral parents have been found, so the origin and evolutionary history of VLEs within fungi is enigmatic.

In recent work, we discovered that a wide range of Saccharomycotina yeasts contain VLEs, including killer plasmids, helper plasmids, and cryptic plasmids (Heneghan et al. 2025a). We also found that many species of Pezizomycotina (filamentous ascomycetes) such as *Fusarium* have killer toxin gene clusters in their nuclear genomes, containing ribonuclease, chitinase, and immunity genes (but not housekeeping genes), which we hypothesized were formed by integration of VLE-like sequence(s) into the nuclear genome, even though free-living VLEs have never been found in Pezizomycotina (Heneghan et al. 2025b). Apart from these gene clusters in Pezizomycotina, we did not find VLEs or VLE-derived killer toxin genes in any other fungal taxa, except for a handful of cases of recent horizontal transfers of toxin gene clusters from Pezizomycotina into Basidiomycota species (Fig. 1A) (Heneghan et al. 2025b). Phylogenetic analysis of chitinases indicated that the chitinase genes in the Pezizomycotina nuclear killer toxin gene clusters and the VLE-encoded chitinases of Saccharomycotina formed sister clades, suggesting that they are orthologs (Heneghan et al. 2025b). This finding raised the possibility that the association of VLEs with fungi might pre-date the origin of Dikarya, and therefore that VLEs might exist in deeper fungal lineages (Fig. 1A). Here, motivated by the recent growth in genome sequence data available from fungi outside Dikarya, and the evidence of infections by *Nucleocytoviricota* (Myers et al. 2025), we searched for VLEs in genome sequence assemblies from non-Dikarya fungi. We report that they are present in both of the other two phyla of terrestrial fungi, Zoopagomycota and Mucoromycota (Mondo and Grigoriev 2025), either as free (presumably cytosolic) VLE molecules or integrated into the nuclear genome. We find that some Zoopagomycota and Mucoromycota species harbor large numbers of VLEs simultaneously. The sequence diversity of DNAPs in non-Dikarya VLEs greatly exceeds the diversity in Saccharomycotina VLEs, and we find that separate clades of killer and helper VLEs were already established by the time of the common ancestor of all terrestrial fungi.

## Results

### VLEs in *Cunninghamella blakesleeana* and other Mucoromycota species

We discovered a set of 7 VLEs in a strain of *Cunninghamella blakesleeana*, a species in phylum Mucoromycota. These VLEs are not present in the reference genome sequence of *C. blakesleeana* (strain AS 3.970, accession number JAFCIZ010000000), which is 32 Mb in size. We found them by accident when we assembled a set of Illumina reads downloaded from NCBI, that had been described as originating from the budding yeast species *Meyerozyma guilliermondii* (strain VKA1, isolated from compost; (Valsalan and Mathew 2020)). The assembly we obtained from these reads was 46 Mb in size, which is much larger than we expected based on the 11 Mb reference assembly of the type strain of *M. guilliermondii* (Butler et al. 2009). By BLASTN analysis we found that the 46 Mb assembly contained contigs from *C. blakesleeana* as well as *M. guilliermondii*, including rDNAs from both species. It consists of contigs that essentially comprise a complete *M. guilliermondii* genome (11 Mb) plus a complete *C. blakesleeana* genome (32 Mb). Therefore, we believe that the original DNA sample used to construct the Illumina sequencing library was of mixed species origin. In the same publication, Valsalan and Mathew (2020) also sequenced the genome of *M. guilliermondii* VKA1 using Nanopore technology and obtained an assembly 11 Mb in size that contains only *M. guilliermondii* rDNA. Since there are no VLEs in this 11 Mb assembly and no VLEs in the assemblies of multiple other *M. guilliermondii* strains available at NCBI, we infer that the VLEs in our 46 Mb assembly come from *C. blakesleeana*.

We identified seven *C. blakesleeana* VLE contigs, ranging in size from 3.9 kb to 14.5 kb (Fig. 2A). Two of them (Node86 and Node87) were initially identified by using chitinase proteins (toxin α/β subunits) from Saccharomycotina VLEs (Heneghan et al. 2025a) as queries in TBLASTN searches against the 46 Mb assembly. Both of these contigs also contain DNAP genes similar to the B-type DNAP genes of Saccharomycotina VLEs. Node86 and Node87 resemble Saccharomycotina killer plasmids. As well as containing 1 or 2 genes for a secreted protein with a chitinase (GH18) domain that is potentially a toxin α/β subunit, they also contain a gene for a small, secreted protein that is potentially a toxin γ-subunit (ribonuclease). The candidate γ subunits from the two contigs have 32% amino acid sequence identity to each other, but no statistically significant similarity to the γ subunits we previously identified in Saccharomycotina and Pezizomycotina species (Heneghan et al. 2025a; Heneghan et al. 2025b). Further TBLASTN searches using the DNAPs from Node86 and Node87 as queries against the 46 Mb assembly led to the identification of five other VLE contigs that also contain DNAP genes (Fig. 2A). One contig, Node83, resembles the helper plasmids seen in Saccharomycotina (Satwika et al. 2012). It contains homologs of all 9 housekeeping genes (named *YKP3* to *YKP11*) that are present in Saccharomycotina helper plasmids, and some of these genes are arranged in the same, highly conserved, order as seen in Saccharomycotina (Heneghan et al. 2025a). A smaller contig, Node109, contains second copies of *YKP5* and *YKP10*, with ∼80% identity to those on Node83. The other VLE contigs in the assembly contain DNAP genes and some ORFs of unknown function (Fig. 2A). One ORF has similarity to the Pep4 protease of *S. cerevisiae*.

**Figure 2.**
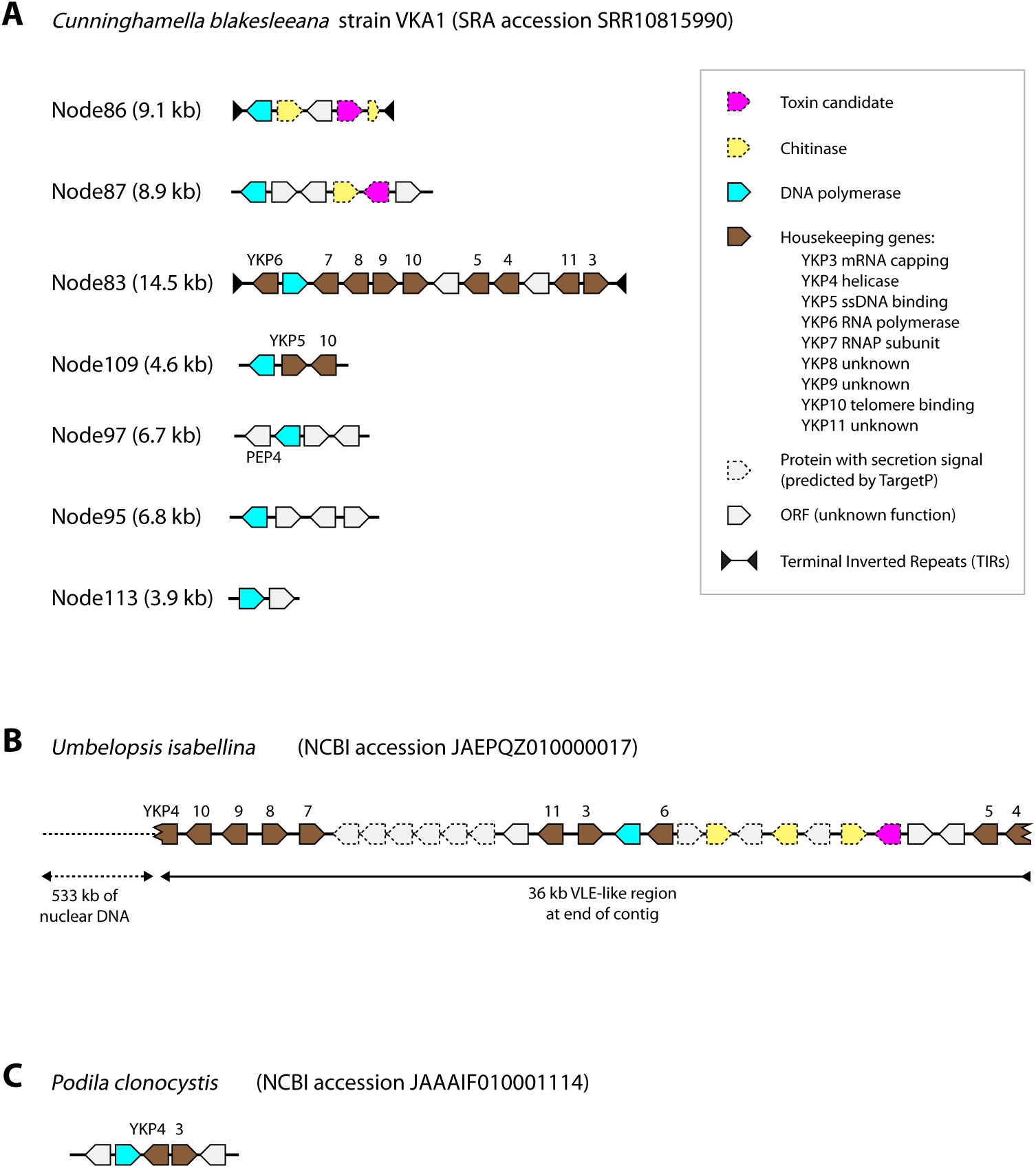
Structures of Mucoromycota VLE-like contigs. (**A**) *Cunninghamella blakesleeana*, high-coverage contigs (nodes) from a *de novo* sequence assembly of isolate VKA1. (B) *Umbelopsis isabellina*, 36 kb VLE-like region at the end of a large nuclear contig in NCBI. (C) *Podila clonocystis*, 10 kb contig in an NCBI genome assembly. Predicted gene functions are as shown in the key.

The killer plasmid-like contig Node86 has terminal inverted repeats (TIRs) at its ends, at least 127 bp long. Similar TIRs of at least 77 bp are present on the helper-like plasmid Node83. TIRs are a characteristic feature of linear dsDNA plasmids replicated by B-type DNAPs, and their presence in Node86 and Node83 indicates that these contigs correspond to free VLEs, i.e. linear dsDNA molecules, rather than parts of the nuclear genome of *C. blakesleeana*. TIRs are not present in the other *C. blakesleeana* VLE-like contigs, but TIRs are often absent from contigs assembled from VLEs because the covalently attached terminal protein at the 5’ ends of the genome interferes with library construction (Heneghan et al. 2025a). All 7*C. blakesleeana* VLE-like contigs have elevated read-coverage in the genome assembly, ranging from 2 to 11 times the coverage of the nuclear genome (Table S1), which suggests that they may all be free VLEs.

We identified VLE-like sequences in two other species in phylum Mucoromycota, by using the *C. blakesleeana* VLE-encoded proteins as queries in TBLASTN searches of genome assemblies in the NCBI databases. First, a contig from *Umbelopsis isabellina*, which is 569 kb long and therefore must be part of a nuclear chromosome, contains a 36-kb VLE-like region at one end (Fig. 2B). The VLE-like region appears to be a complete fused killer-helper plasmid. It includes three chitinase genes, a DNAP gene, and a complete set of nine housekeeping genes (*YKP3* to *YKP11*). Its *YKP4*, the helicase gene, is broken into two parts that are located at the ends of the VLE-like region and have a perfect 127-bp overlap. It also contains 10 genes for predicted secreted proteins, as well as three ORFs without secretion signals. One of the secreted proteins is a candidate *Umbelopsis* γ-toxin because it has sequence similarity to the two toxin candidates we identified in *Cunninghamella*: 72% identity to the Node87 candidate, and 29% identity to the Node86 candidate (Fig. 3). Second, we found a small VLE-like contig in a genome assembly of *Podila clonocystis*, containing a DNAP gene, *YKP3*, *YKP4* and two unidentified ORFs, but no γ-toxin candidate (Fig. 2C). The read coverage of this contig is 60x that of the nuclear genome (Table S1). *Podila* is a genus in the Mucoromycota subphylum Mortierellomycotina, whereas *Cunninghamella* and *Umbelopsis* are in subphylum Mucoromycotina (Fig. 1A).

**Figure 3.**
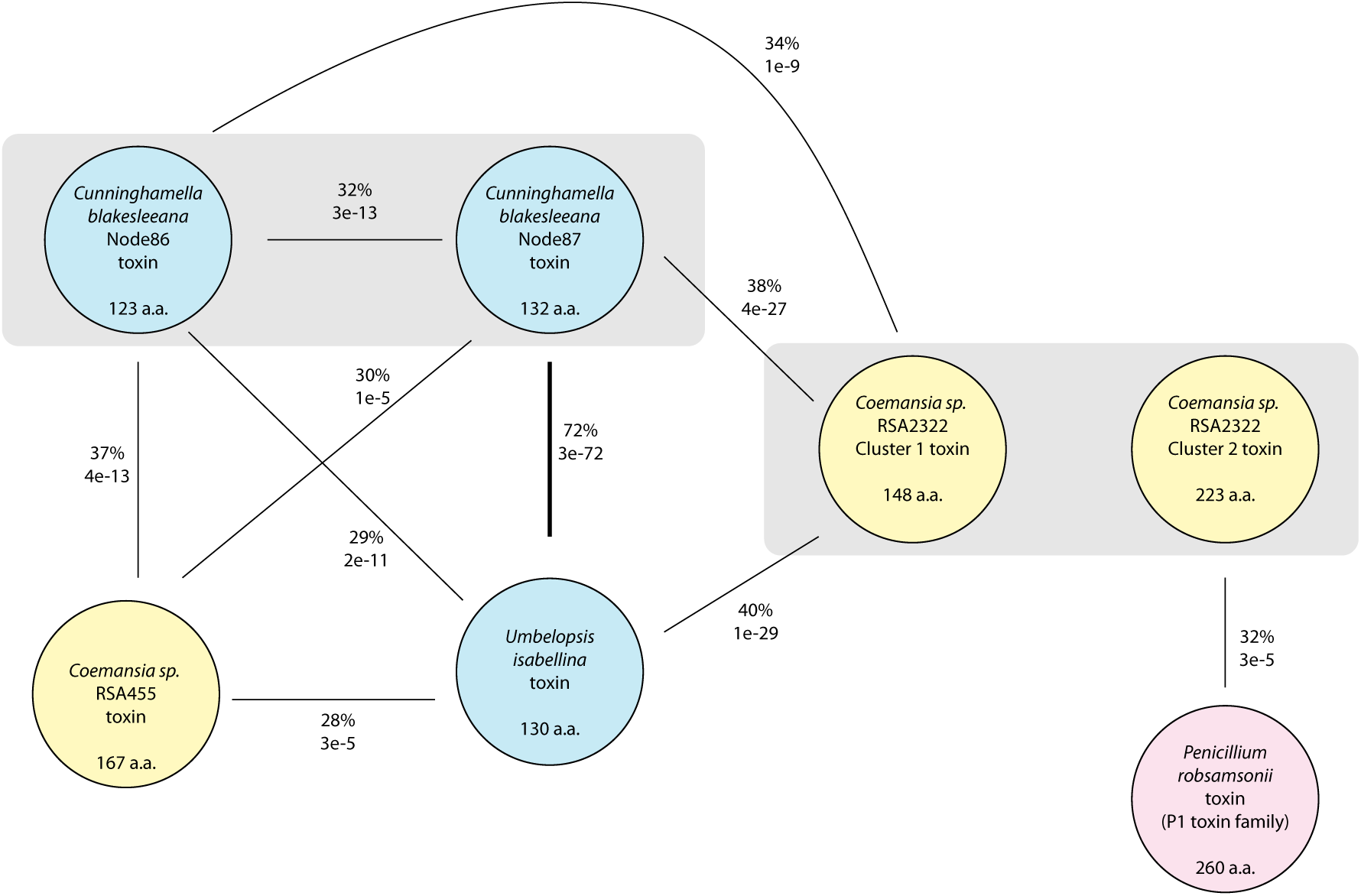
Sequence relationships among putative γ-toxins encoded by non-Dikarya VLEs. Circles represent toxin candidates, colored by phylum (blue, Mucoromycota; yellow, Zoopagomycota; red, Ascomycota). Connecting lines indicate sequence similarity, showing the BLASTP E-value and the percent amino acid sequence identity in the region aligned by BLASTP. Absence of a connecting line indicates absence of a BLASTP hit (E-value weaker than 1e-4). Gray boxes indicate toxin candidates made by the same organism. Protein lengths are shown.

### Constellations of VLEs in *Coemansia* species (Zoopagomycota)

We discovered that *Coemansia*, a genus in phylum Zoopagomycota, contains large numbers of VLEs (Fig. 4). We initially detected these VLEs when we carried out TBLASTN searches using DNAP, chitinase, or housekeeping (*YKP*) genes as queries, against all fungal genome assemblies (excluding Dikarya) in the NCBI Whole Genome Shotgun database. These searches returned large numbers of significant hits to contigs from different *Coemansia* isolates, particularly when DNAP sequences were used as queries (Table S2). *Coemansia* is a genus in the fungal family Kickxellaceae, in subphylum Kickxellomycotina of phylum Zoopagomycota (Fig. 1A). A substantial effort to investigate the genomes and phylogeny of this family was made by Reynolds *et al*. (2023), who used low-coverage Illumina genome sequencing to sequence 171 Kickxellaceae species including more than 100 *Coemansia* isolates. Many of the *Coemansia* isolates represent new, unnamed species and are identified only by their strain numbers. Reynolds *et al*. (2023) showed that the current genus *Coemansia* is polyphyletic, and they found that there are four major clades in family Kickxellaceae, which they referred to as clades *Coemansia* (i.e. the clade containing the type species of the genus), Kickxellales1, Kickxellales2, and *Kickxella*. The number of VLE-like contigs varies considerably among *Coemansia* isolates, and some isolates have few of them, but we detected isolates with ≥6 VLE-like contigs in all four clades of the family, with the highest numbers being in Kickxellales2 (Table S2).

**Figure 4.**
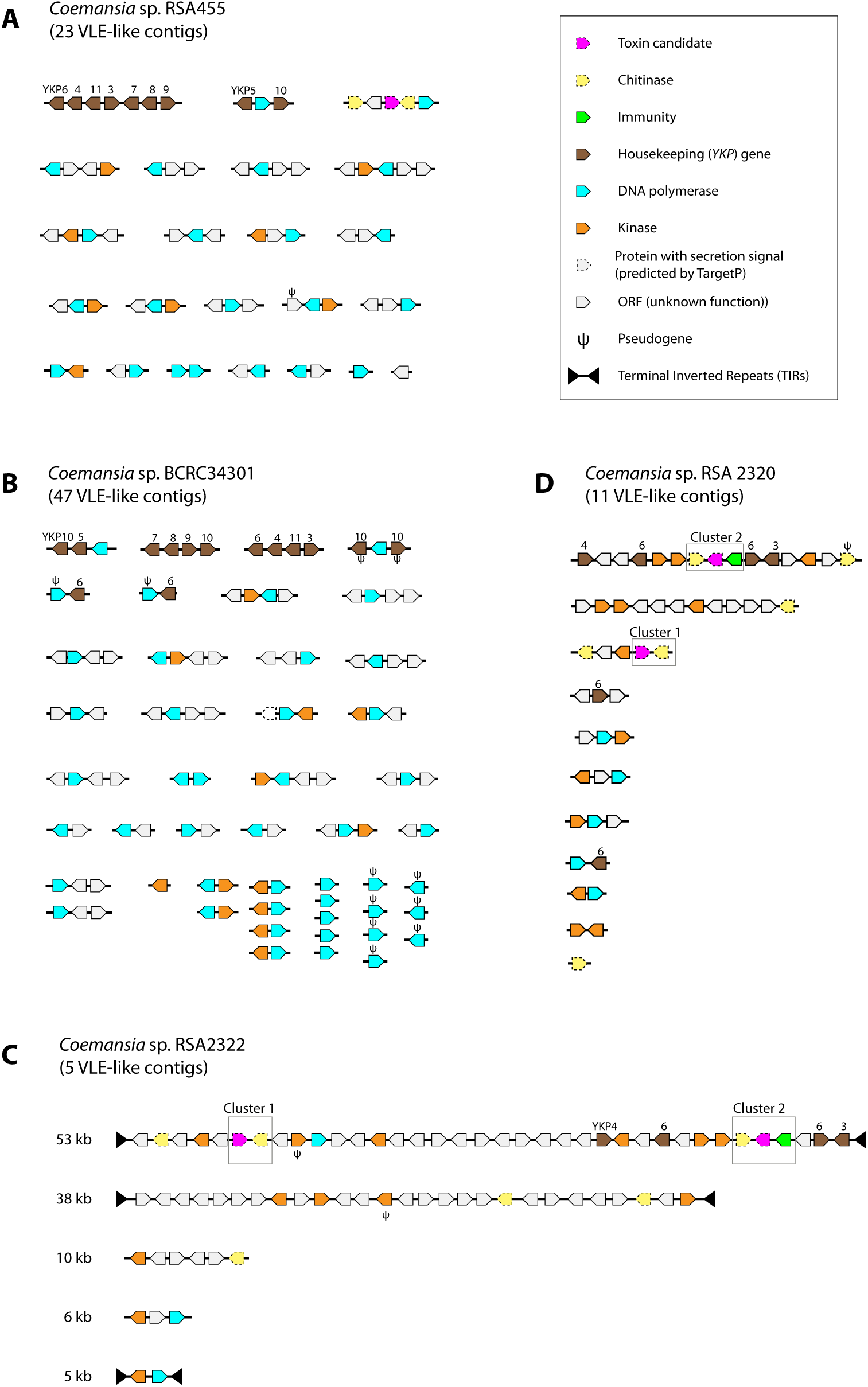
Structures of Zoopagomycota VLE-like contigs identified in *de novo* assemblies of four *Coemansia* species. (A) *Coemansia* sp. RSA455. (B) *Coemansia* sp. BCRC34301. (C) *Coemansia* sp. RSA2322. (D) *Coemansia* sp. RSA2320. Predicted gene functions are as shown in the key.

To further characterize *Coemansia* VLEs, we reassembled the genomes of four isolates of interest, using the Illumina reads deposited in the NCBI SRA database by Reynolds *et al*. (2023). We used the SPAdes assembler (Bankevich et al. 2012) so that we could compare the approximate read coverage of each VLE-like contig to the read coverage of the nuclear genome. All the VLE-like contigs in these isolates have high read coverage (Table S1), and several have TIRs, so we infer that they are derived from free VLEs, not integrated into the nuclear genome.

*Coemansia* sp. RSA455 has 23 VLE contigs (Fig. 4A), whose average read coverage is 22x that of the nuclear genome (range 9x–58x; Table S1). Remarkably, 21 of these VLE contigs contain DNAP genes (Fig. 4A). Because B-family DNA polymerases use the terminal protein bound to the end of the VLE dsDNA as a primer for initiating replication, it seems probable that each VLE replicates independently, using the DNAP polymerase that it encodes. Thus RSA455 appears to maintain 21 orthogonal DNA replication systems. Two RSA455 VLE contigs contain a complete set of housekeeping genes (*YKP3* to *YKP11*) between them. Another VLE contig contains two genes for secreted chitinase-like proteins, similar to the α/β subunit of zymocin, and a gene for a small secreted protein that we considered to be a candidate γ-toxin. The *Coemansia* sp. RSA455 toxin candidate has weak but statistically significant sequence similarity to the toxin candidates from Mucoromycotina (Fig. 3). The other 20 VLE contigs in RSA455 contain several ORFs of unknown function. They include a diverse family of 8 ORFs that are all predicted to code for proteins with a kinase domain.

*Coemansia* sp. BCRC34301 has an even larger constellation of VLEs than RSA455 (Fig. 4B). It has 47 VLE contigs, with read coverage averaging 29x the nuclear genome coverage (range 4x-80x), and 44 of them contain a DNAP gene. In total BCRC34301 has 45 DNAP genes, although 9 of them appear to be pseudogenes with internal stop codons or frameshifts. It also has 13 diverse ORFs with kinase domains, and a complete set of housekeeping (*YKP*) genes including some duplicates. This *Coemansia* isolate does not have any VLE contigs coding for chitinases or a putative γ-toxin, so it is not obvious what selective pressure has maintained its VLEs.

*Coemansia* sp. RSA2322 is of interest because it appears to code for two different toxins, one of which originated by horizontal gene transfer between fungal phyla. It has five VLE-like contigs, with average read coverage 16x that of the nuclear genome, three of which are complete VLEs with TIRs (Fig. 4C). One VLE is 53 kb long, making it the largest VLE known, and it contains two candidate γ-toxin gene clusters. Cluster 1 contains genes for a chitinase and a candidate γ-toxin with highest similarity to those in the Mucomycota species *Cunninghamella* and *Umbelopsis* (Fig. 3). Cluster 2 contains a candidate γ-toxin, chitinase, and putative immunity protein, and resembles (both in sequence and in structure) the 3-gene clusters that we previously identified in Pezizomycotina nuclear genomes (Heneghan et al. 2025b). The RSA2322 Cluster 2 γ-toxin candidate has highest sequence similarity to the γ-toxin from the Pezizomycotina species *Penicillium robsamsonii* (Fig. 3). The two candidate γ-toxins of *Coemansia* sp. RSA2322 do not have any significant sequence similarity to each other, nor to the candidate γ-toxin of *Coemansia* sp. RSA455. Interestingly, RSA2322 does not appear to have a complete set of housekeeping genes; we detected only *YKP3* (mRNA capping enzyme), *YKP4* (helicase) and *YKP6* (RNA polymerase, 2 copies), making it unclear how the VLEs are maintained. Its VLEs also contain several ORFs predicted to be kinases. A fourth *Coemansia* isolate, RSA2320 has VLE contigs that are similar in sequence to those of RSA2322 (and these two isolates are very closely related; (Reynolds et al. 2023)), but RSA2320 has more contigs and more DNAP genes than RSA2322 (Fig. 4D).

To test their functionality, we expressed the genes for three candidate γ-toxins in *S. cerevisiae*, using the same method we previously used to validate toxins from Saccharomycotina and Pezizomycotina species (Heneghan et al. 2025a; Heneghan et al. 2025b). The toxin candidates we tested were from *Coemansia* sp. RSA455, *Coemansia* sp. RSA2322 (cluster 1), and *Umbelopsis isabellina*. Disappointingly, none of these genes caused a growth defect upon induction in *S. cerevisiae* (Fig. S1). This negative result indicates that the genes do not code for toxins that act against *S. cerevisiae*, even though they code for proteins with secretion signals and are located beside chitinase genes, similarly to known γ-toxin genes. It remains possible that these genes code for *bona fide* toxins, but that *S. cerevisiae* is too distantly related to whatever species is their natural target.

### Non-Dikarya VLEs encode killer toxin-like chitinases

The chitinase subunits of zymocin-like killer toxins produced by Saccharomycotina and Pezizomycotina are all members of one phylogenetic subgroup of fungal chitinases, called the C-II subgroup (Tzelepis and Karlsson 2019). They contain a GH18 glycosyl hydrolase domain and chitin-binding domain(s). To investigate how the chitinase genes present in the Mucoromycota and Zoopagomycota VLE contigs are related to other chitinases, we constructed a maximum-likelihood phylogenetic tree using the GH18 domain, which is universally present in fungal chitinases. The phylogenetic tree confirms that all the Mucoromycota/Zoopagomycota VLE-encoded chitinases are members of the C-II subgroup (Fig. 5). It also shows that most of the Mucoromycota/Zoopagomycota chitinases form a clade that is a sister group to the Saccharomycotina and Pezizomycotina killer toxin chitinases, which is concordant with the expected phylogeny of these three phyla/subphyla and suggests that these three large groups of chitinases are orthologs, descended vertically from an ancestor in an early terrestrial fungus.

**Figure 5.**
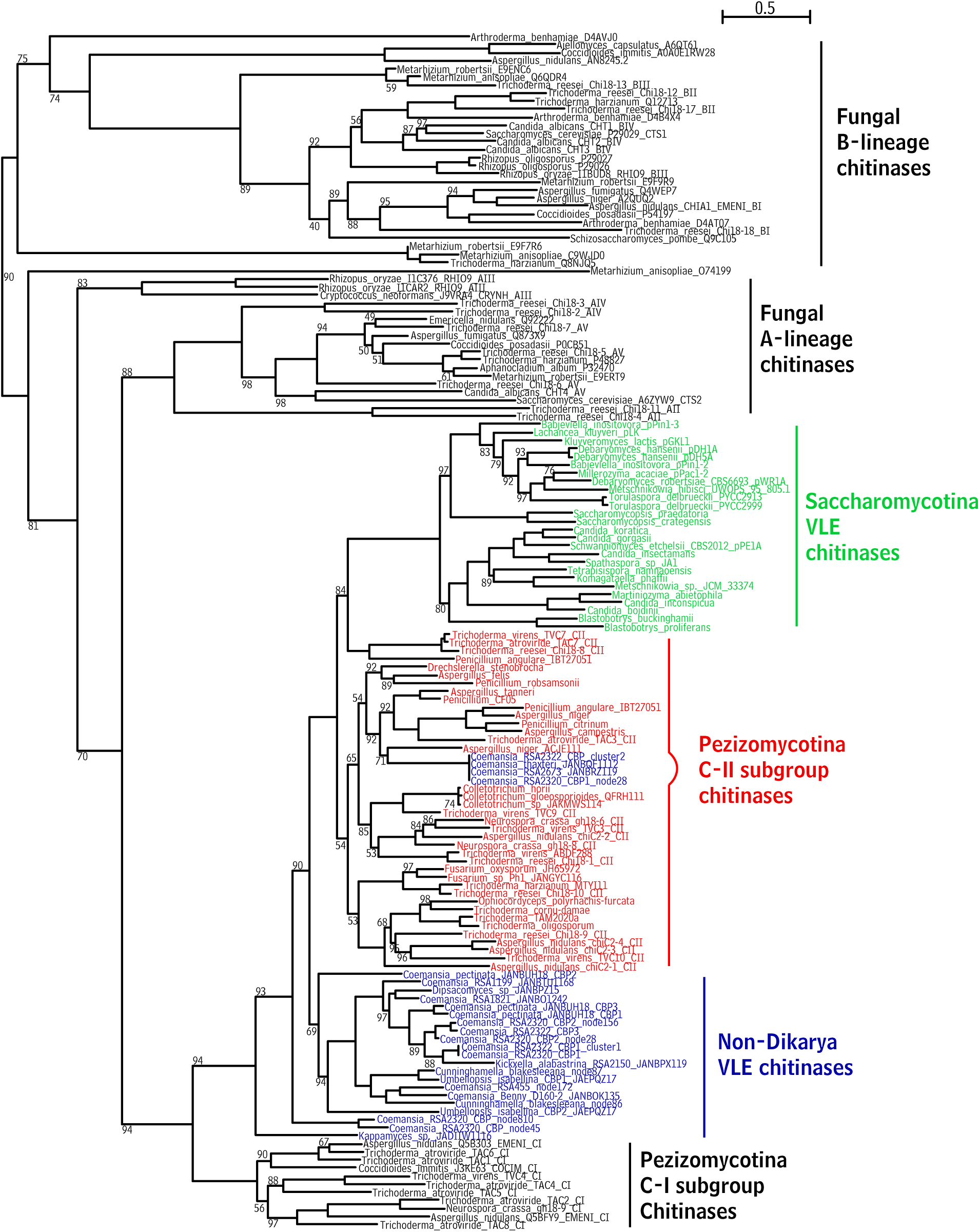
Phylogenetic tree of GH18 domains of fungal chitinase proteins. Chitinase genes located on VLE contigs of non-Dikarya species (Zoopagomycota, Mucoromycota and *Kappamyces*) are named in blue. Chitinase genes from Saccharomycotina VLEs are named in green, and chitinases from Pezizomycotina nuclear toxin gene clusters are named in red. The tree was constructed using maximum likelihood methods and rooted using fungal A– and B-lineage chitinase sequences. All bootstrap values below 99% are shown; unlabeled branches have 100% support.

Strikingly, a group of four *Coemansia* chitinases is placed at a position within the main Pezizomycotina clade, quite separate from all the other Mucoromycota/Zoopagomycota chitinases (Fig. 5). This group includes the cluster 2 chitinase gene of *Coemansia* sp. RSA2322, as well as highly similar genes from VLEs in three other *Coemansia* isolates (RSA2320, RSA2673 and *Coemansia thaxteri*). Each of these C-II chitinases is located beside a gene with high similarity to Pezizomycotina immunity genes, and a γ-toxin candidate with highest similarity to Pezizomycotina γ-toxins. None of the other Mucoromycota/Zoopagomycota VLEs contain identifiable immunity genes. Thus, all three genes in toxin gene cluster 2 from RSA2322 appear to have originated by inter-phylum horizontal gene transfer from a Pezizomycotina donor to a Zoopagomycota (*Coemansia*) recipient, presumably in a single event that transferred a complete three-gene cluster. In RSA2322, the horizontally transferred genes became integrated into a cytoplasmic element rather than into the nuclear genome, and they integrated into a VLE that already contained another γ-toxin gene cluster (cluster 1). For the three other similar *Coemansia* isolates, each of the assemblies has homologs of cluster 1 and cluster 2 on separate high-copy contigs that may be different VLEs.

By TBLASTN searches of the NCBI databases, we also identified a single VLE-like contig in a *Kappamyces* species (Fig. S2). *Kappamyces* is a member of Chytridiomycota, a phylum of zoosporic fungi (Fig. 1A). The VLE-like contig is in *Kappamyces* sp. JEL0829, which is one of only two isolates of this genus whose genomes have been sequenced (Amses et al. 2022). The contig is 26 kb long and contains genes for a C-II chitinase and two immunity proteins beside a 221 amino acid secreted protein that is a candidate γ-toxin, as well as containing the conserved VLE housekeeping genes *YKP6* (RNA polymerase), *YKP4* (helicase) and *YKP9* (unknown function), and several unidentified ORFs (Fig. S2). We do not have information about the read coverage of this contig. Phylogenetic analysis of the chitinase places it at a position in the C-II clade outside all the Zoopagomycota, Mucoromycota and Ascomycota C-II chitinases (Fig. 5), consistent with the phylogenetic position expected for Chytridiomycota. The candidate γ-toxin has no significant homologs in database searches. Although this contig suggests that killer plasmid-like VLEs exist in Chytridiomycota, the possibility of contamination cannot be ruled out and the conclusion must remain tentative until further examples of Chytridiomycota VLEs are found.

### Non-Dikarya VLE DNAPs form 5 phylogenetic groups

There is high sequence divergence among the VLE DNAPs in the *Coemansia* species. To investigate how the DNAPs and, by extension, the VLEs are related to one another, we constructed a phylogenetic tree using all the near full-length DNAP sequences encoded by VLEs from six *Coemansia* species that have multiple DNAPs: three species discussed above (RSA455, BCRC34301, and RSA2320, which are in the Kickxellales2 clade and have 22, 39, and 5 near full-length DNAPs, respectively), two other Kickxellales2 species BCRC34181 (13 DNAPs) and S146 (18 DNAPs), and a Kickxellales1 species RSA2049 (12 DNAPs). The DNAPs are highly divergent in sequence from each other. They form five distinct phylogenetic groups (Fig. 6), and it is evident that the diversity of VLEs in *Coemansia* is due to amplifications that occurred in four of these groups (Groups 1-4). Amplification appears to be ongoing, because different species of *Coemansia* show different levels of amplification in different DNAP groups. For example, the high number of DNAPs in BCRC34301 is mostly due to the presence of many VLEs with Group 1 DNAPs, whereas most of the DNAPs in RSA2049 are in Groups 3 and 4.

**Figure 6.**
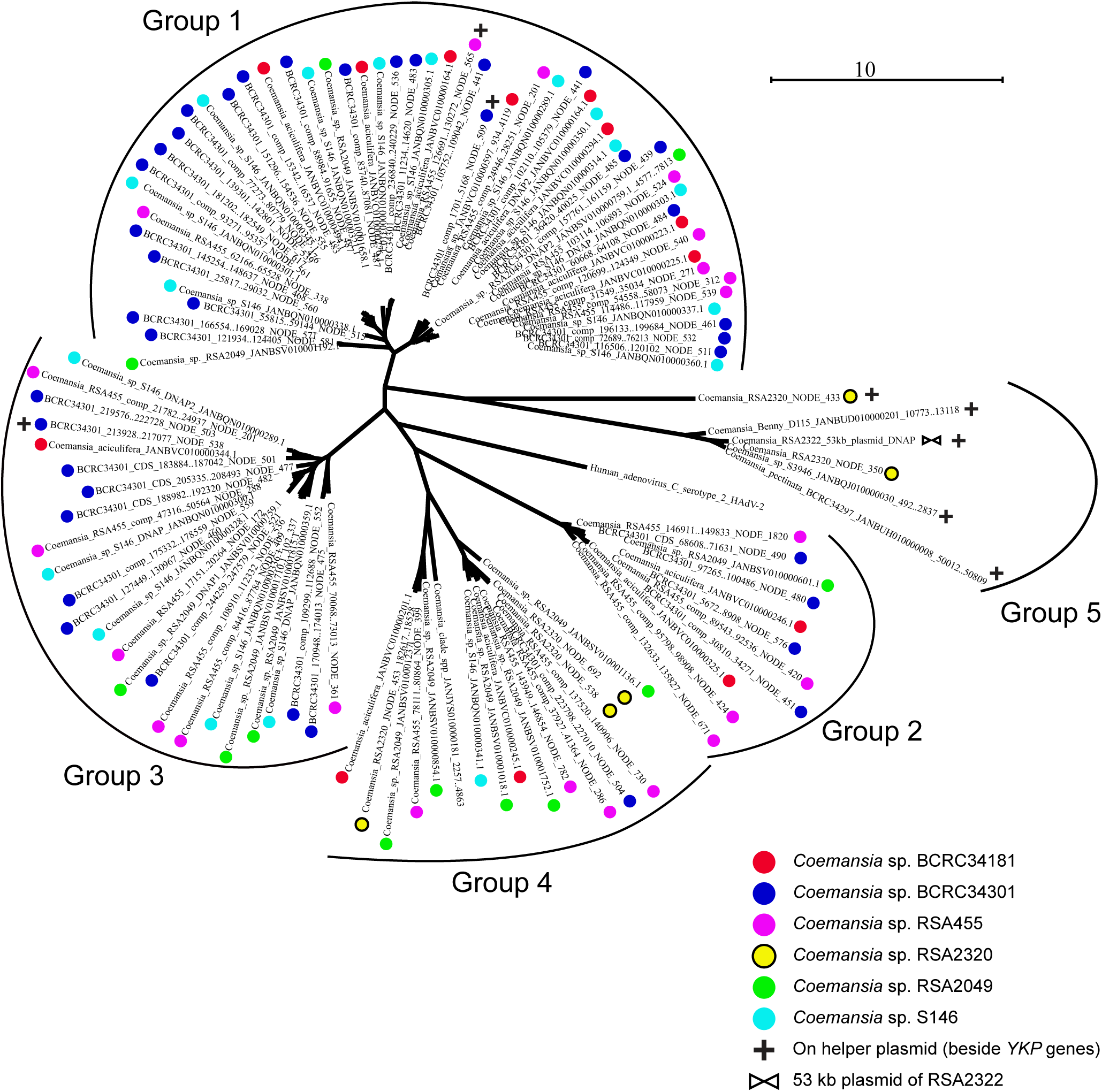
DNA polymerases in *Coemansia* species VLEs form five phylogenetic groups. The phylogenetic tree shows the relationships among the DNAP (pPolB) proteins from 6 different *Coemansia* species that each have multiple VLE-like contigs containing DNAP genes. Colored dots indicate the different species. DNAPs from some other *Coemansia* species that only have a single VLE DNAP were included and fall into Group 5, which also contains the DNAP present on the 53 kb TIR-containing VLE from *Coemansia sp.* RSA2322. DNAP of a human adenovirus was included as a potential outgroup but the tree is shown in unrooted format. Cross symbols mark DNAP genes that are located on contigs that also contain YKP (housekeeping) genes.

RSA2320, which is an outgroup to the other Kickxellelales2 species examined (Reynolds et al. 2023), only has DNAPs belonging to Groups 4 and 5 (Fig. 6).

To relate the Mucoromycota DNAPs to the five groups of DNAPs that we identified in *Coemansia* (Zoopagomycota), we constructed a phylogenetic tree that included Mucoromycota DNAPs and one representative DNAP per group from each *Coemansia* species (Fig. S3). Five of the seven DNAPs on the *Cunninghamella blakesleena* VLEs fall into Group 4, including the DNAPs from its two killer-like VLEs (Node86 and Node87; Fig. 2A). The DNAP of its helper plasmid-like VLE (Node83) falls into Group 5, as do the DNAPs from helper-like VLE contigs from *Umbelopsis* and *Podila*. It is notable that in *Coemansia*, Group 5 also contains DNAPs from several helper-like VLEs (e.g. RSA2320 Node433, which contains *YKP6* and a DNAP) or helper-killer fusion VLEs (e.g. the 53 kb VLE of RSA2322) and it has not undergone the same extent of amplification as other groups (Fig. 6). The seventh DNAP of *Cunninghamella blakesleeana* (Node97) falls in Group 1. Overall, it appears that the divergence among most of the groups of DNAPs, and therefore among the groups of VLEs that contain them, predated the divergence between the phyla Zoopagomycota and Mucoromycota, because both of these phyla maintain DNAPs from Groups 1, 4 and 5.

### An ancient divergence between killer and helper VLEs in terrestrial fungi

We investigated how the DNAPs of Saccharomycotina VLEs are related to the five phylogenetic groups of DNAPs we identified in VLEs of non-Dikarya fungi (Fig. 7). In Saccharomycotina, all helper VLEs contain a DNAP gene, whereas only some killer and cryptic VLEs contain one; the others rely on helper-encoded DNAPs (Heneghan et al. 2025a). We found that the Saccharomycotina DNAPs are most closely related to non-Dikarya Groups 1 and 3, so the diversity of VLE DNAPs in Saccharomycotina is only a small subset of the diversity in non-Dikarya species. All the Saccharomycotina killer VLE DNAPs cluster with non-Dikarya Group 3, which forms a close sister group to them (Fig. 7). Each of the DNAPs from Saccharomycotina helper VLEs clusters with either Group 1 or Group 3. The Saccharomycotina VLE DNAPs are distant from non-Dikarya Groups 2 and 4. Non-Dikarya Group 5, which contains *Coemansia* and most of the Mucoromycota DNAPs, lies outside a large clade (96% bootstrap support) that contains DNAPs from many Saccharomycotina helper VLEs and also contains non-Dikarya Group 1, which consists of several *Coemansia* DNAPs and one DNAP from *Cunninghamella*.

**Figure 7.**
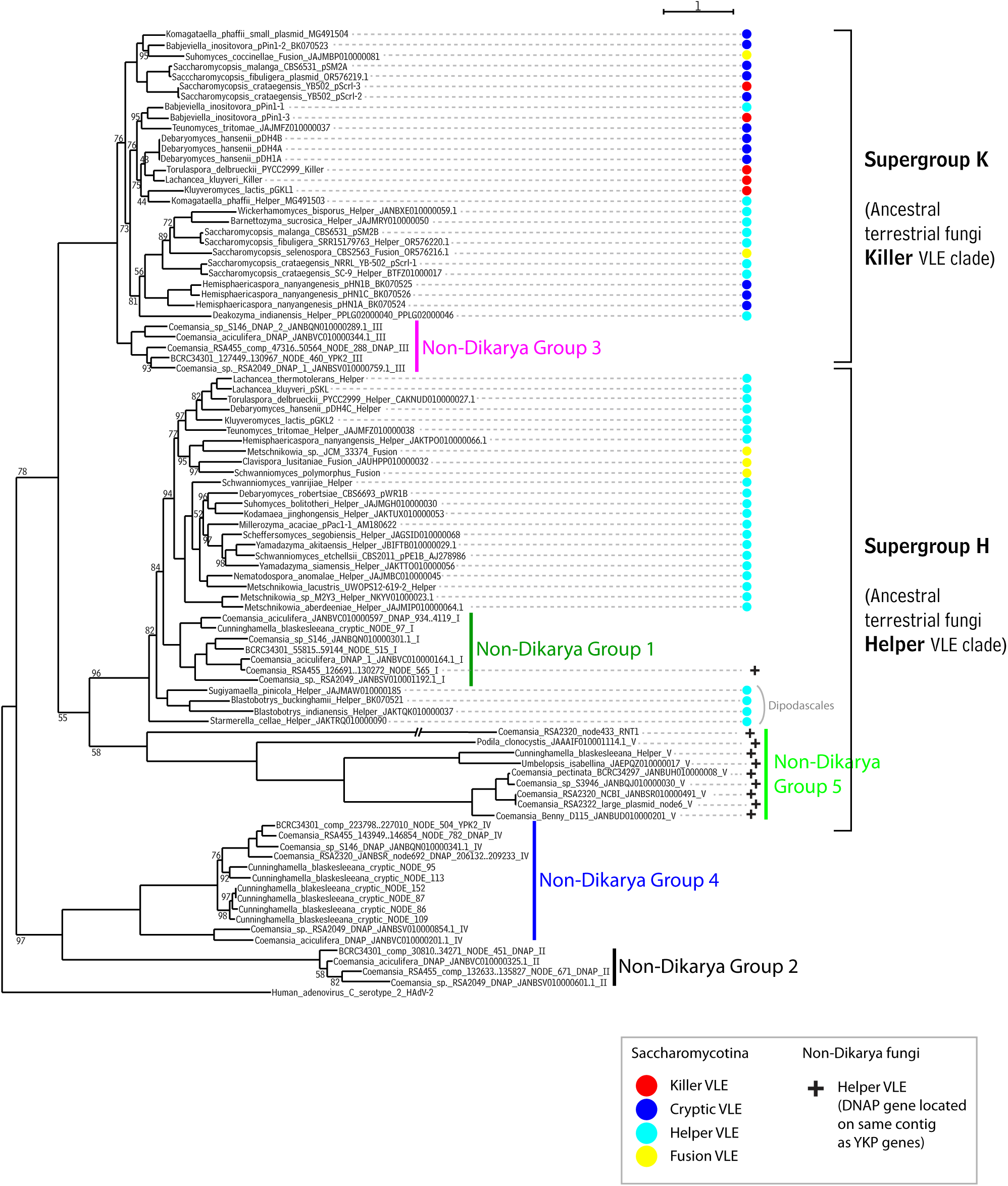
Phylogeny of DNA polymerases suggests that the division between killer and helper VLEs is ancient. DNAP proteins from Saccharomycotina VLEs were compared to DNAPs from Zoopagomycota and Mucoromycota VLE contigs. Saccharomycotina VLEs are classified as killer, helper, cryptic, or fusion (fused killer-helper) plasmids as in the key. The 5 phylogenetic groups of Zoopagomycota/Mucoromycota DNAPs identified in Fig. 6 are labeled. Cross symbols indicate Zoopagomycota/Mucoromycota DNAPs that are located on probable helper VLEs, i.e. VLE contigs that also contain *YKP* housekeeping genes. The tree was constructed by maximum likelihood using Modelfinder and IQ-Tree, bootstrapped for 1000 replicates. Bootstrap values below 100% are shown. The branch for one DNAP in Clade 5 (the DNAP from Coemansia sp. RSA2320 NODE433, which unusually contains an RNT1 ribonuclease domain as well as a DNAP domain) is drawn at 50% of its true length to save space.

Examination of the DNAP phylogenetic tree suggests that the distinction between killer and helper VLEs is an ancient divergence that has been present since the origin of terrestrial fungi. We propose that there are ancestral killer and helper VLE clades as shown in Fig. 7 and described further below.

The ancestral killer clade, which we refer to as Supergroup K, includes all the Saccharomycotina killer VLE DNAPs, and the non-Dikarya Group 3 DNAPs (Fig. 7). The DNAPs of all Saccharomycotina cryptic VLEs also lie in this clade. Cryptic VLEs are VLEs that lack housekeeping (*YKP*) genes but do not carry an identifiable γ-toxin gene, although in many cases they contain a chitinase (toxin α/β subunit) gene (Polomska et al. 2021; Heneghan et al. 2025a). The DNAP tree strongly suggests that cryptic VLEs are derivatives of killer VLEs. Supergroup K also contains DNAPs from some helper VLEs, but we suggest that these are the result of relatively recent gene exchange events in which a helper VLE replaced its original DNAP gene with a DNAP gene from the killer VLE present in the same species. An example of this process can be seen in *Babjeviella inositovora*, whose helper VLE (pPin1-1) and killer VLE (pPin1-3) have DNAP genes with very similar sequences, both of which lie in Supergroup K.

The ancestral helper clade, which we refer to as Supergroup H, consists of DNAPs from the majority of Saccharomycotina helper VLEs, as well as non-Dikarya Groups 1 and 5 (Fig. 7). There are no Saccharomycotina killer or cryptic VLEs in this clade. We suggest that Group 5 is the non-Dikarya sister clade to Saccharomycotina helper VLE clade, because it includes the DNAPs of the helper VLEs (i.e. VLEs that also contain *YKP* housekeeping genes) of three Mucoromycota species as well as *Coemansia* species. Interestingly, all the helper VLE DNAPs from species of Dipodascales, which is the most deeply divergent order within Saccharomycotina that contains VLEs (Heneghan et al. 2025a), lie at a position in Supergroup H outside a clade containing non-Dikarya Group 1 and helper DNAPs from all the other orders of Saccharomycotina (Fig. 7). This topology suggests that non-Dikarya Group 1 arose by horizontal transfer from Saccharomycotina to Zoopagomycota, after the orders of Saccharomycotina had begun to diverge. Alternatively, it is possible that the tree topology in Fig. 7 is slightly wrong and that Group 1 is actually the non-Dikarya sister clade to the Saccharomycotina helper VLE clade, rather than being nested inside it; bootstrap support values for the two critical branches are 82% and 84%. Regardless of which of these alternatives is correct, we can conclude that there were two separate supergroups of DNAPs, one associated with helper VLEs and one associated with killer VLEs, in the common ancestor of terrestrial fungi.

Lastly, fusion VLEs, which are large dsDNAs that were formed by fusion between a helper VLE and a killer or cryptic VLE, can contain a DNAP from either Supergroup K or Supergroup H (Fig. 7). From a functional point of view, because the DNAP interacts with the TIRs at each end of the VLE, it makes sense that a fusion VLE should retain only one DNAP gene and that it could come from either clade.

## Discussion

We have found the first evidence that homologs of yeast virus-like elements (VLEs) exist in terrestrial fungal taxa outside the ascomycete subphylum Saccharomycotina. Our searches uncovered clear evidence of free VLEs, in high copy number and separate from the nuclear genome, in the genera *Coemansia* (Zoopagomycota) and *Cunninghamella* (Mucoromycota), as well as evidence of VLEs integrated into the nuclear genome in some other Mucoromycota species. We did not find any evidence of VLEs in any zoosporic fungal phyla, except for a single example of a possible VLE in *Kappamyces*. Similar to the situation in Saccharomycotina, it appears that VLEs are present in only a small percentage of species, because the databases we searched included all the whole-genome shotgun assemblies available from non-Dikarya fungi in the current NCBI databases, and the VLEs we report here are the only ones we found. Even in the genus *Coemansia*, we found that high numbers of VLEs are rare (e.g., only 17 of the 161 available assemblies had ≥10 contigs containing DNAPs; Table S2). Another similarity to Saccharomycotina is that VLEs can be present in some isolates of a species but absent in others; we found this to be the case in *Cunninghamella blakesleeana*.

Some of the Mucoromycota and Zoopagomycota VLEs contain ORFs coding for secreted proteins, reminiscent of killer γ-toxins like zymocin, and they are located beside chitinase genes that could be used as toxin α/β subunits. However, our laboratory experiments with three of these non-Dikarya toxin candidates failed to confirm this hypothesis because they did not cause a growth defect in *Saccharomyces cerevisiae*. There are several possible explanations for the lack of toxicity, including that they may be optimized for a target in Mucoromycota or Zoopagomycota that does not exist in *S. cerevisiae*. It is not even clear whether the toxin candidates could potentially be anticodon nucleases – the proteins are significantly shorter than in Ascomycota (range 130-167 amino acid residues in Mucoromycota and Zoopagomycota, compared to 225-351 in Saccharomycotina and 226-366 in Pezizomycotina), and downstream of the secretion signal, most do not have the Glu-8 residue that is well conserved among Saccharomycotina and Pezizomycotina γ-toxins (Meineke et al. 2012; Chakravarty et al. 2014; Heneghan et al. 2025b). They could possibly be an entirely different cargo delivered by a chitinase. Furthermore, the presence in some *Coemansia* species (e.g. *Coemansia* sp. BCRC34301; Fig. 4B) of many VLEs that do not code for either a chitinase, a secreted γ-toxin candidate, or a *YKP* housekeeping gene, suggests that these VLEs must be maintained by selection for the production of a different protein product, possibly a kinase or the product of an unidentified ORF. Most puzzlingly, the set of 47 VLE-like contigs in BCRC34301 codes for only one predicted secreted protein, an ORF of 138 amino acids that has no homologs in sequence databases, not even in other *Coemansia* species.

Our phylogenetic analysis of DNAP sequences indicated that a separation of VLEs into killer types and helper types is ancient. A possible advantage of this arrangement is that it may provide modularity, allowing one toxin to be replaced by another, perhaps to target a different victim, while retaining a set of helper genes that are already well adapted to the host. It could also enable multiple killer VLEs, coding for different toxins, to be maintained in the same cell by a single helper VLE.

Although we did see some evidence of inter-phylum horizontal gene transfer – most notably the transfer of a complete toxin gene cluster from Pezizomycotina to *Coemansia* sp. RSA2322 – the predominant picture that emerges from our phylogenetic analyses of chitinases, killer VLE DNAPs, and helper VLE DNAPs is one of vertical inheritance and a sister relationship (orthology) between VLE genes of Saccharomycotina and VLE genes of non-Dikarya fungi. Our results suggest the model of fungal VLE evolution shown in Fig. 8. The ancestor of terrestrial fungi contained multiple VLEs that were already differentiated into at least four types: a killer type (Supergroup K), a helper type (Supergroup H), and two unknown types (Groups 2 and 4). These four types are still present in Zoopagomycota. After the speciation that separated the ancestor of Zoopagomycota from the ancestor of Saccharomycotina, Saccharomycotina retained only a subset of the previous VLE diversity: a killer VLE with a DNAP from Supergroup K, and a helper VLE with a DNAP from Supergroup H. The Saccharomycotina helper VLEs subsequently diverged in sequence as the orders of Saccharomycotina separated. At some point after Dipodascales had diverged from the other Saccharomycotina orders, a helper VLE was transferred horizontally from Saccharomycotina to Zoopagomycota, resulting in the non-Dikarya clade Group 1. Other Zoopagomycota retained their original helper VLE, resulting in the non-Dikarya clade Group 5. There may also have been a horizontal transfer of a Group 1 VLE from Zoopagomycota to Mucoromycota, which would explain the presence of both a Group 1 DNAP and a Group 5 DNAP in *Cunninghamella*.

**Figure 8.**
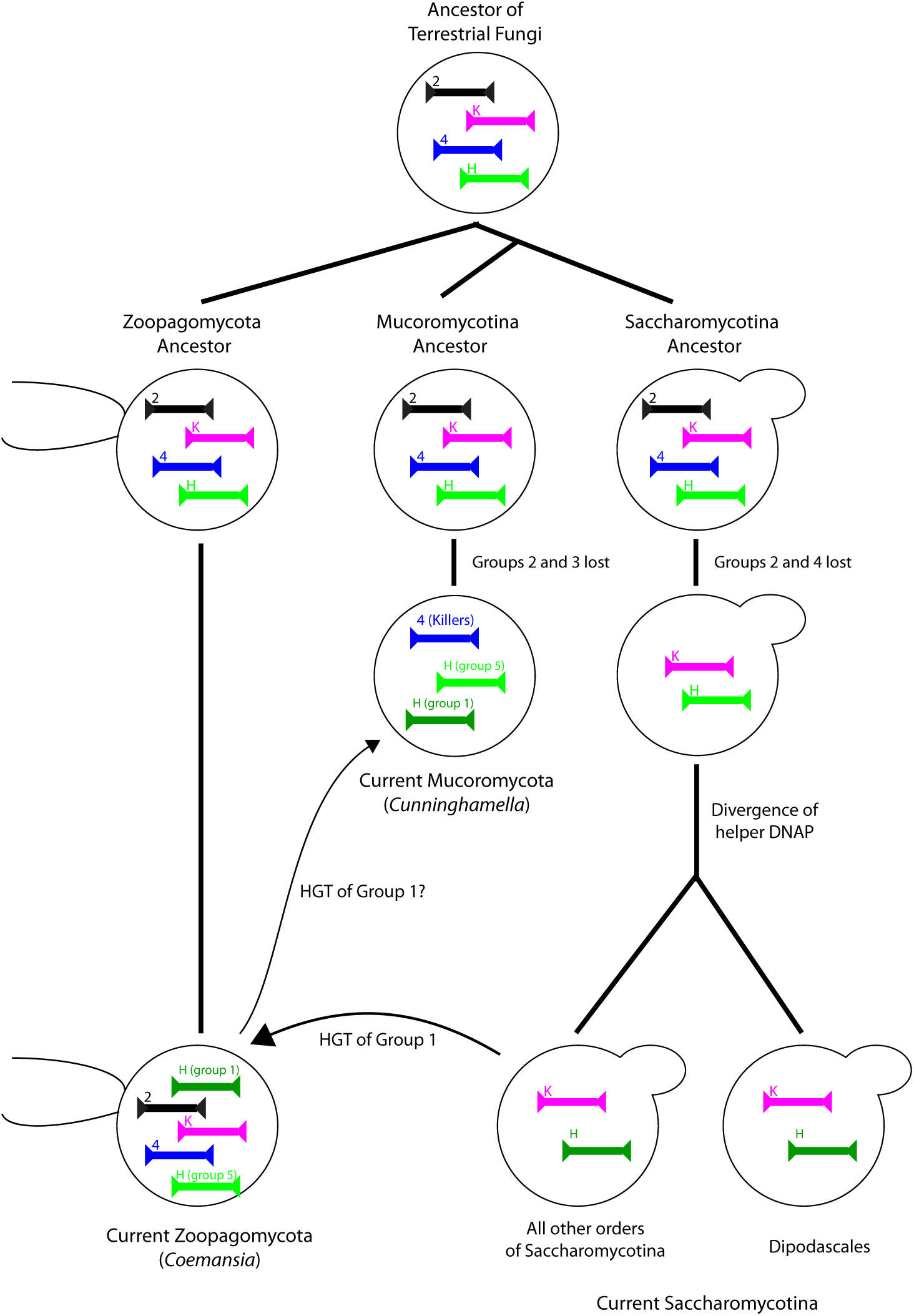
Hypothesis for evolution of VLEs in terrestrial fungi. See Discussion. Labels 2 and 4 denote DNAP phylogenetic groups 2 and 4. Labels K and H denote Supergroup K (killer, which includes group 3) and Supergroup H (helper, which includes groups 1 and 5) respectively.

Lastly, we note that non-Dikarya VLEs may have some biotechnological applications. If the multi-VLE systems of Zoopagomycota and Mucoromycota contain genes coding for toxins, they may be antifungal agents with novel mechanisms of action, so characterizing their activity and targets should be a priority. It is also intriguing that dozens of linear dsDNAs, each apparently replicating independently by using its own DNAP, can coexist within a *Coemansia* cell. In principle, they could be used to develop multiply orthogonal synthetic replication systems for rapid laboratory evolution of multiple chosen target sequences, similar but more extensive than those developed from the two *Kluyveromyces lactis* VLEs (Arzumanyan et al. 2018; Ravikumar et al. 2018). In addition, *Cunninghamella* has potential to be developed as a model system for testing antiviral inhibition of *Nucleocytoviricota* such as poxviruses (Sykora et al. 2018).

## Methods

The VLEs and VLE-like contigs in non-Dikarya species reported here were identified from TBLASTN searches of the NCBI whole-genome shotgun sequence database, taxonomically restricted to Fungi excluding Dikarya, using amino acid sequences of chitinase, DNAP, or YKP3-YKP11 proteins from Saccharomycotina VLEs as queries. The Saccharomycotina VLEs were described in our previous study (Heneghan et al. 2025a). For four *Coemansia* species (RSA455, BCRC34301, RSA2322 and RSA2320), we initially identified VLEs in the existing assemblies (Reynolds et al. 2023) but we also made *de novo* assemblies from Illumina reads available in the NCBI SRA database, in order to estimate the read-coverage of their VLE contigs relative to their nuclear contigs (Table S1). We assembled NCBI SRA Illumina reads (accession SRR10815990) from the isolate VKA1 (Valsalan and Mathew 2020), which we found to be a mixture of reads from *Cunninghamella blakesleeana* and *Meyerozyma guilliermondii*. Illumina reads were assembled using SPAdes version 3.14 (Bankevich et al. 2012). We also assembled reads (SRR5192073) from *Podila clonocystis*.

To estimate the approximate numbers of DNAP genes in genome assemblies from Zoopagomycota species (Table S2), we carried out a series of TBLASTN searches using the NCBI BLAST webserver. The database was all Zoopagomycota genome assemblies in the Whole Genome Shotgun database (Sept. 2025). As queries, we used all the VLE DNAP proteins we had previously manually annotated from *Coemansia* species RSA455, RSA2320, and RSA2322, all DNAPs from *Cunninghamella blakesleeana* VLEs, and the DNAP from *Babjeviella inositovora* pPin1-2. This set of queries includes representatives of each of the non-Dikarya DNAP clades 1-5. All contigs that were hit with a TBLASTN E-value *E* < 1e-10 were retained, and the results were parsed to count the number of unique contigs that were hit in each species.

Secretion signals were predicted using the TargetP webserver (https://services.healthtech.dtu.dk/services/TargetP-2.0/), using a cutoff value of 0.95 (Emanuelsson et al. 2007). For phylogenetic analysis, protein multiple sequence alignments were made using MUSCLE (Edgar 2004), and trees were constructed using IQ-Tree with model selection by ModelFinder (Minh et al. 2020). Expression of γ-toxin candidate genes in *Saccharomyces cerevisiae* was carried using the same β-estradiol induction system as described previously (Heneghan et al. 2025a; Heneghan et al. 2025b).

## Acknowledgments

This work was supported by the European Research Council (789341).

## Notes

### Competing Interest Statement

The authors have declared no competing interest.

