## Supplementary Information for "Zymocin-like killer plasmids were present in the common ancestor of terrestrial fungi"

Figures S1-S3.

Tables S1 and S2.

**Figure S1**. Induction of expression of γ-toxin candidates from Zoopagomycota and Mucoromycota does not affect growth of *Saccharomyces cerevisiae*. Candidate toxin genes, without their predicted secretion signals, were synthesized and cloned into *S. cerevisiae* under the control of a promoter inducible by the hormone β-estradiol, as in (Heneghan et al. 2025a). Growth curves over 24 hours are shown for the toxin constructs in the presence and absence of the inducer, and for a control (empty vector) construct. No significant reduction of growth of *S. cerevisiae* is seen after induction. The candidate toxin genes tested were from (A) *Coemansia* sp. RSA2322 cluster 1 (bases 7601..7996 of NCBI accession number JANBSQ010000019.1; strain PHY090); (B) *Coemansia* sp. RSA455 (bases 4005..4454 of NCBI accession number JANBRM010000335.1; strain PHY126); (C) *Umbelopsis isabellina* (bases 565512..565174 of NCBI accession number JAEPQZ010000017.1; strain PHY089).

**Figure S2.** A VLE-like contig from the Chytridiomycota species *Kappamyces* sp. JEL0829 (NCBI accession number JADIIW010000116). The chitinase gene codes for a C-II chitinase whose phylogenetic position is shown in Fig. 5. The candidate γ-toxin and the chitinase both contain a Cys residue near their C-terminus, consistent with formation of a disulfide bond between them as in zymocin. The two candidate immunity proteins Imm1 and Imm2 have 43% amino acid sequence identity to each other. In database searches they have highest similarity to a candidate immunity protein from *Coemansia* sp. RSA2320 (BLASTP scores 6e-27 and 41% identity for Imm1; 7e-26 and 31% identity for Imm2). They also have significant sequence identity to candidate immunity proteins from Pezizomycotina (e.g. BLASTP score 8e-22 and 35% identity for Imm2 versus *Aspergillus felis*). Not drawn to scale. White boxes indicate ORFs of unknown function. Brown boxes indicate housekeeping (*YKP*) genes with homologs on Saccharomycotina helper VLEs. The contig does not have TIRs.

**Figure S3.** Unrooted phylogenetic tree of VLE DNAPs from Mucoromycota and Zoopagomycota. A maximum likelihood tree was made using Modelfinder in IQ-Tree and bootstrapped for 1000 replicates. Colored dots indicate different *Coemansia* or Mucoromycota species as indicated in the legend. Only one sequence from each *Coemansia* species was included from each group identified in Fig. 6. The long branch from *Coemansia* sp. RSA2320 corresponds to a DNAP that unusually has a RNT1-like domain, a ribonuclease.

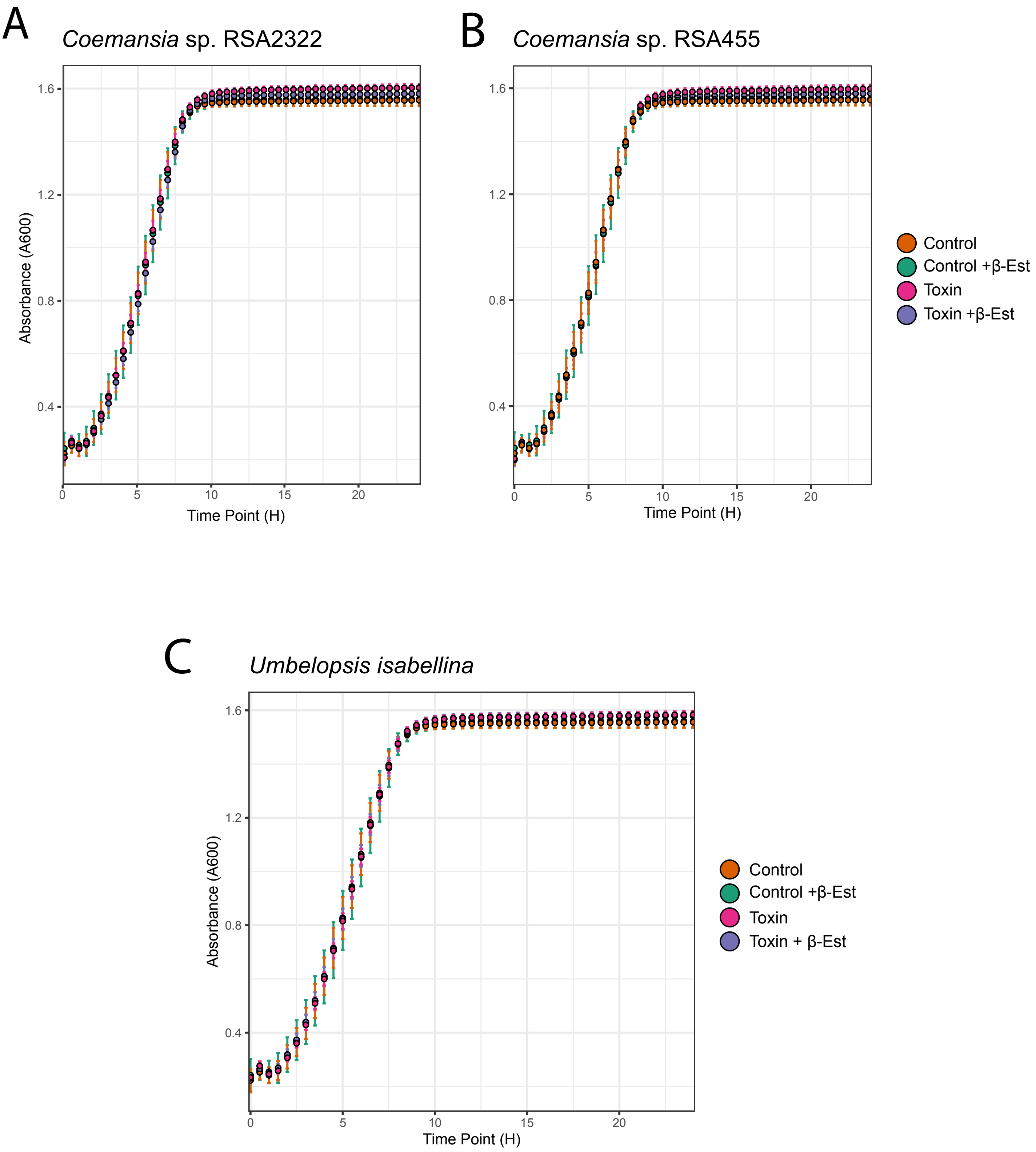
A
Coemansia sp. RSA2322
B
Coemansia sp. RSA455
Absorbance (A600)
Time Point (H)
1.6
1.2
0.8
0.4
0
5
10
15
20
Control
Control +β-Est
Toxin
Toxin +β-Est
C
Umbelopsis isabellina
Absorbance (A600)
Time Point (H)
Control
Control +β-Est
Toxin
Toxin + β-Est

Figure S1

*Kappamyces* sp. JEL0829 VLE-like contig.

NCBI accession number JADIIW010000116.   26294 bp.

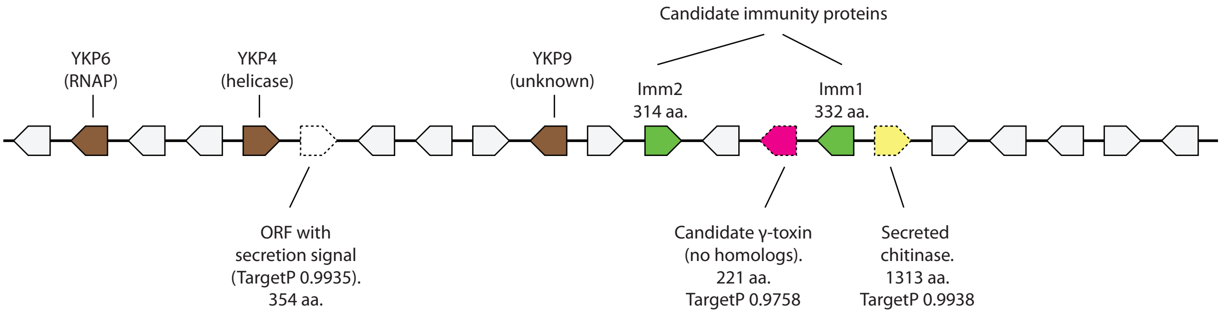

Figure S2

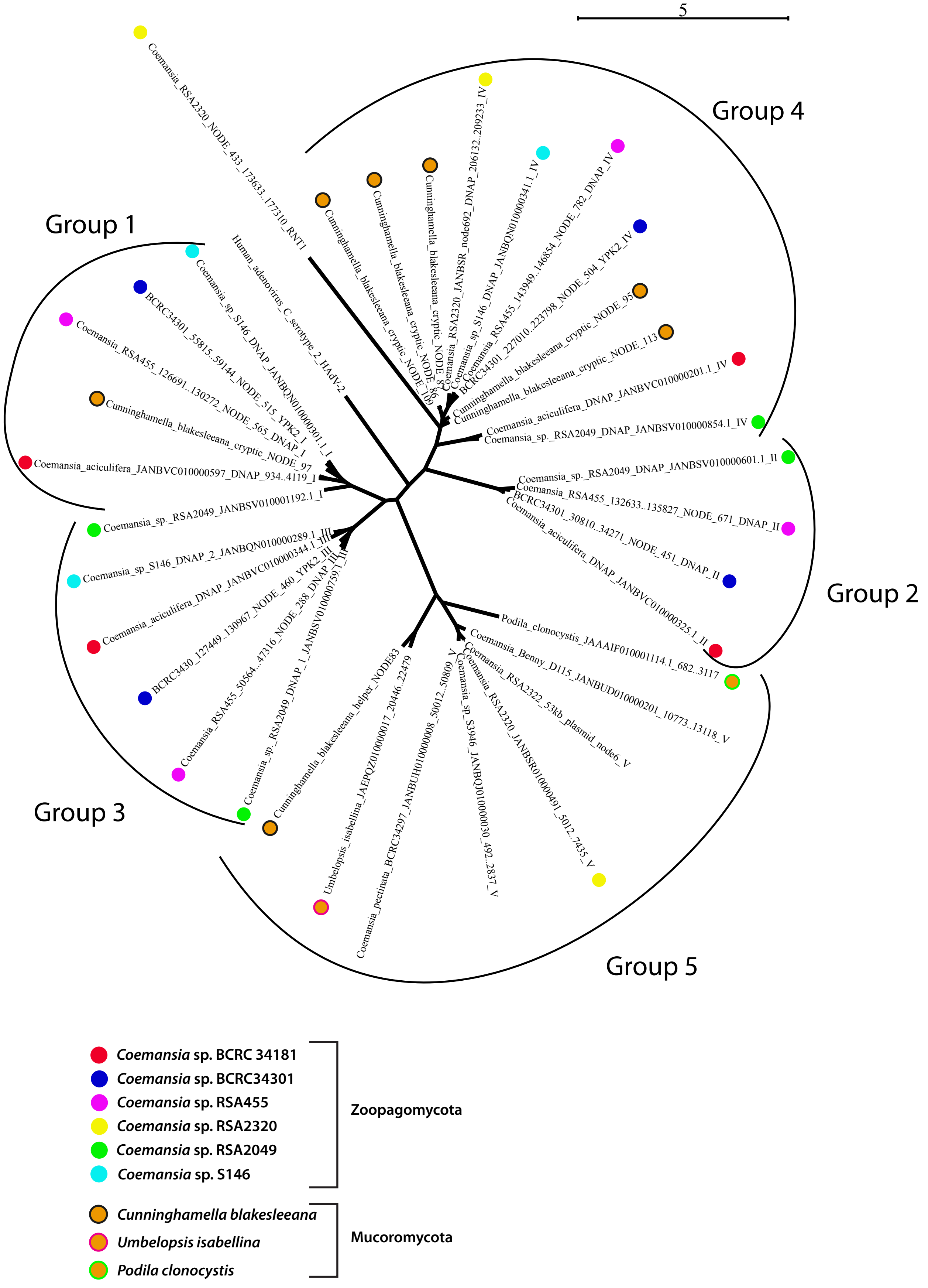
5
Group 1
Group 2
Group 3
Group 4
Group 5
Coemansia_RSA2320_NODE_433_173633..177310_RNT1
Human_adenovirus_C_serotype_2_HAdV-2
Coemansia_sp_S146_DNAP_JANBQN010000301.1_I
BCRC34301_55815..59144_NODE_515_YPK2_I
Coemansia_RSA455_126691..130272_NODE_565_DNAP_I
Cunninghamella_blakesleeana_cryptic_NODE_97
Coemansia_aciculifera_JANBVC010000597_DNAP_934..4119_I
Coemansia_sp._RSA2049_JANBSV010001192.1_I
Coemansia_sp_S146_DNAP_2_JANBQN010000289.1_III
Coemansia_aciculifera_DNAP_JANBVC010000344.1_III
BCRC3430_127449..130967_NODE_460_YPK2_III
Coemansia_RSA455_50564..47316_NODE_288_DNAP_III
Coemansia_sp._RSA2049_DNAP_1_JANBSV010000759.1_III
Cunninghamella_blakesleeana_helper_NODE83
Umbelopsis_isabellina_JAEPQZ010000017_20446..22479
Coemansia_pectinata_BCRC34297_JANBUH010000008_50012..50809_V
Coemansia_sp_S3946_JANBQJ010000030_492..2837_V
Coemansia_RSA2320_JANBSR010000491_5012..7435_V
Coemansia_RSA2322_53kb_plasmid_node6_V
Coemansia_Benny_D115_JANBUD010000201_10773..13118_V
Podila_clonocystis_JAAAIF010001114.1_682..3117
Coemansia_aciculifera_DNAP_JANBVC010000325.1_II
BCRC34301_30810..34271_NODE_451_DNAP_II
Coemansia_RSA455_132633..135827_NODE_671_DNAP_II
Coemansia_sp._RSA2049_DNAP_JANBSV010000601.1_II
Coemansia_sp._RSA2049_DNAP_JANBSV010000854.1_IV
Coemansia_aciculifera_DNAP_JANBVC010000201.1_IV
Cunninghamella_blakesleeana_cryptic_NODE_113
Cunninghamella_blakesleeana_cryptic_NODE_95
BCRC34301_227010..223798_NODE_504_YPK2_IV
Coemansia_RSA455_143949..146854_NODE_782_DNAP_IV
Coemansia_sp_S146_DNAP_JANBQN010000341.1_IV
Coemansia_RSA2320_JANBSR_node692_DNAP_206132..209233_IV
Cunninghamella_blakesleeana_cryptic_NODE_86
Cunninghamella_blakesleeana_cryptic_NODE_109
Coemansia sp. BCRC 34181
Coemansia sp. BCRC34301
Coemansia sp. RSA455
Coemansia sp. RSA2320
Coemansia sp. RSA2049
Coemansia sp. S146
Zoopagomycota
Cunninghamella blakesleeana
Umbelopsis isabellina
Podila clonocystis
Mucoromycota

Figure S3

**Table S1. Read coverage in *de novo* assemblies made with the SPAdes assembler.**

| Species (SRA No.) | Node number | Length (bp) | Absolute coverage | Coverage relative to nuclear genome | Mean relative coverage of VLE-like contigs |
|---|---|---|---|---|---|
| **Cunninghamella blakesleeana (SRR10815990)** | **93** | **6910** | **17.9** | **1.0** | |
| Cunninghamella blakesleeana (SRR10815990) | 83 | 14563 | 197.0 | 11.0 | |
| Cunninghamella blakesleeana (SRR10815990) | 86 | 9107 | 84.9 | 4.7 | |
| Cunninghamella blakesleeana (SRR10815990) | 87 | 8897 | 103.5 | 5.8 | |
| Cunninghamella blakesleeana (SRR10815990) | 95 | 6799 | 35.0 | 2.0 | |
| Cunninghamella blakesleeana (SRR10815990) | 97 | 6688 | 126.6 | 7.1 | |
| Cunninghamella blakesleeana (SRR10815990) | 109 | 4629 | 166.8 | 9.3 | |
| Cunninghamella blakesleeana (SRR10815990) | 113 | 3916 | 127.0 | 7.1 | 6.7 |
| **Podila clonocystis (SRR5192073)** | **1** | **175417** | **4.6** | **1.0** | |
| Podila clonocystis (SRR5192073) | 957 | 10544 | 275.6 | 59.9 | 59.9 |
| **Coemansia sp. RSA455 (SRR10999133)** | **1** | **153469** | **12.9** | **1.0** | |
| Coemansia sp. RSA455 (SRR10999133) | 148 | 10472 | 550.9 | 42.7 | |
| Coemansia sp. RSA455 (SRR10999133) | 172 | 9818 | 330.8 | 25.6 | |
| Coemansia sp. RSA455 (SRR10999133) | 201 | 9183 | 396.0 | 30.7 | |
| Coemansia sp. RSA455 (SRR10999133) | 271 | 7933 | 146.6 | 11.4 | |
| Coemansia sp. RSA455 (SRR10999133) | 286 | 7754 | 123.8 | 9.6 | |
| Coemansia sp. RSA455 (SRR10999133) | 288 | 7735 | 231.9 | 18.0 | |
| Coemansia sp. RSA455 (SRR10999133) | 312 | 7501 | 245.5 | 19.0 | |
| Coemansia sp. RSA455 (SRR10999133) | 338 | 7246 | 756.2 | 58.6 | |
| Coemansia sp. RSA455 (SRR10999133) | 361 | 7042 | 131.7 | 10.2 | |
| Coemansia sp. RSA455 (SRR10999133) | 399 | 6763 | 183.8 | 14.2 | |
| Coemansia sp. RSA455 (SRR10999133) | 409 | 6716 | 312.4 | 24.2 | |
| Coemansia sp. RSA455 (SRR10999133) | 420 | 6637 | 336.1 | 26.1 | |
| Coemansia sp. RSA455 (SRR10999133) | 424 | 6550 | 281.8 | 21.8 | |
| Coemansia sp. RSA455 (SRR10999133) | 524 | 6109 | 396.1 | 30.7 | |
| Coemansia sp. RSA455 (SRR10999133) | 536 | 6064 | 195.2 | 15.1 | |
| Coemansia sp. RSA455 (SRR10999133) | 539 | 6064 | 195.2 | 15.1 | |
| Coemansia sp. RSA455 (SRR10999133) | 540 | 6044 | 227.5 | 17.6 | |
| Coemansia sp. RSA455 (SRR10999133) | 565 | 5885 | 754.8 | 58.5 | |
| Coemansia sp. RSA455 (SRR10999133) | 671 | 5359 | 204.8 | 15.9 | |
| Coemansia sp. RSA455 (SRR10999133) | 730 | 5087 | 185.3 | 14.4 | |
| Coemansia sp. RSA455 (SRR10999133) | 782 | 4903 | 125.1 | 9.7 | |
| Coemansia sp. RSA455 (SRR10999133) | 1820 | 2979 | 161.1 | 12.5 | |
| Coemansia sp. RSA455 (SRR10999133) | 5602 | 1111 | 182.8 | 14.2 | 22.4 |
| **Coemansia sp. BCRC34301 (SRR10999308)** | **1** | **253076** | **14.3** | **1.0** | |
| Coemansia sp. BCRC34301 (SRR10999308) | 429 | 8198 | 626.1 | 43.8 | |
| Coemansia sp. BCRC34301 (SRR10999308) | 439 | 7622 | 242.5 | 17.0 | |
| Coemansia sp. BCRC34301 (SRR10999308) | 441 | 7544 | 395.3 | 27.6 | |
| Coemansia sp. BCRC34301 (SRR10999308) | 451 | 7194 | 702.5 | 49.1 | |
| Coemansia sp. BCRC34301 (SRR10999308) | 460 | 6885 | 325.3 | 22.7 | |
| Coemansia sp. BCRC34301 (SRR10999308) | 461 | 6861 | 172.8 | 12.1 | |
| Coemansia sp. BCRC34301 (SRR10999308) | 468 | 6759 | 258.5 | 18.1 | |
| Coemansia sp. BCRC34301 (SRR10999308) | 475 | 6296 | 212.4 | 14.9 | |
| Coemansia sp. BCRC34301 (SRR10999308) | 476 | 6270 | 499.6 | 34.9 | |
| Coemansia sp. BCRC34301 (SRR10999308) | 477 | 6270 | 153.7 | 10.7 | |
| Coemansia sp. BCRC34301 (SRR10999308) | 480 | 6164 | 414.0 | 29.0 | |
| Coemansia sp. BCRC34301 (SRR10999308) | 482 | 6042 | 175.2 | 12.3 | |
| Coemansia sp. BCRC34301 (SRR10999308) | 483 | 6034 | 845.0 | 59.1 | |
| Coemansia sp. BCRC34301 (SRR10999308) | 484 | 6034 | 845.0 | 59.1 | |
| Coemansia sp. BCRC34301 (SRR10999308) | 485 | 5994 | 677.0 | 47.3 | |
| Coemansia sp. BCRC34301 (SRR10999308) | 487 | 5933 | 473.4 | 33.1 | |
| Coemansia sp. BCRC34301 (SRR10999308) | 490 | 5873 | 524.0 | 36.6 | |
| Coemansia sp. BCRC34301 (SRR10999308) | 496 | 5759 | 778.2 | 54.4 | |
| Coemansia sp. BCRC34301 (SRR10999308) | 501 | 5650 | 176.4 | 12.3 | |
| Coemansia sp. BCRC34301 (SRR10999308) | 503 | 5573 | 124.3 | 8.7 | |
| Coemansia sp. BCRC34301 (SRR10999308) | 504 | 5569 | 121.9 | 8.5 | |
| Coemansia sp. BCRC34301 (SRR10999308) | 509 | 5415 | 1142.4 | 79.9 | |
| Coemansia sp. BCRC34301 (SRR10999308) | 511 | 5378 | 362.4 | 25.3 | |
| Coemansia sp. BCRC34301 (SRR10999308) | 515 | 5229 | 587.5 | 41.1 | |
| Coemansia sp. BCRC34301 (SRR10999308) | 521 | 5059 | 61.5 | 4.3 | |
| Coemansia sp. BCRC34301 (SRR10999308) | 528 | 4937 | 313.7 | 21.9 | |
| Coemansia sp. BCRC34301 (SRR10999308) | 532 | 4845 | 505.3 | 35.3 | |
| Coemansia sp. BCRC34301 (SRR10999308) | 536 | 4729 | 67.6 | 4.7 | |
| Coemansia sp. BCRC34301 (SRR10999308) | 538 | 4706 | 136.3 | 9.5 | |
| Coemansia sp. BCRC34301 (SRR10999308) | 539 | 4644 | 266.3 | 18.6 | |

| | | | | | |
|---|---|---|---|---|---|
| Coemansia sp. BCRC34301 (SRR10999308) | 552 | 4484 | 392.2 | 27.4 | |
| Coemansia sp. BCRC34301 (SRR10999308) | 555 | 4372 | 247.1 | 17.3 | |
| Coemansia sp. BCRC34301 (SRR10999308) | 559 | 4312 | 201.2 | 14.1 | |
| Coemansia sp. BCRC34301 (SRR10999308) | 560 | 4309 | 710.7 | 49.7 | |
| Coemansia sp. BCRC34301 (SRR10999308) | 561 | 4287 | 438.8 | 30.7 | |
| Coemansia sp. BCRC34301 (SRR10999308) | 571 | 3963 | 240.2 | 16.8 | |
| Coemansia sp. BCRC34301 (SRR10999308) | 576 | 3892 | 881.0 | 61.6 | |
| Coemansia sp. BCRC34301 (SRR10999308) | 581 | 3734 | 341.6 | 23.9 | |
| Coemansia sp. BCRC34301 (SRR10999308) | 600 | 3253 | 595.8 | 41.7 | |
| Coemansia sp. BCRC34301 (SRR10999308) | 617 | 2912 | 197.6 | 13.8 | |
| Coemansia sp. BCRC34301 (SRR10999308) | 679 | 2231 | 430.3 | 30.1 | |
| Coemansia sp. BCRC34301 (SRR10999308) | 704 | 1946 | 824.5 | 57.7 | |
| Coemansia sp. BCRC34301 (SRR10999308) | 728 | 1723 | 240.3 | 16.8 | |
| Coemansia sp. BCRC34301 (SRR10999308) | 737 | 1677 | 726.4 | 50.8 | |
| Coemansia sp. BCRC34301 (SRR10999308) | 768 | 1481 | 105.0 | 7.3 | |
| Coemansia sp. BCRC34301 (SRR10999308) | 853 | 1185 | 245.3 | 17.2 | |
| Coemansia sp. BCRC34301 (SRR10999308) | 865 | 1166 | 622.7 | 43.5 | 29.2 |
| **Coemansia sp. RSA2322 (SRR10999316)** | **1** | **241433** | **10.7** | **1.0** | |
| Coemansia sp. RSA2322 (SRR10999316) | 6 | 53497 | 69.8 | 6.5 | |
| Coemansia sp. RSA2322 (SRR10999316) | 7 | 38410 | 62.9 | 5.9 | |
| Coemansia sp. RSA2322 (SRR10999316) | 27 | 10093 | 216.2 | 20.2 | |
| Coemansia sp. RSA2322 (SRR10999316) | 43 | 6173 | 420.4 | 39.3 | |
| Coemansia sp. RSA2322 (SRR10999316) | 50 | 4581 | 108.5 | 10.1 | 16.4 |
| **Coemansia sp. RSA2320 (SRS4604247)** | **1** | **131105** | **52.6** | **1.0** | |
| Coemansia sp. RSA2320 (SRS4604247) | 28 | 38594 | 548.3 | 10.4 | |
| Coemansia sp. RSA2320 (SRS4604247) | 45 | 31633 | 415.1 | 7.9 | |
| Coemansia sp. RSA2320 (SRS4604247) | 156 | 15759 | 469.2 | 8.9 | |
| Coemansia sp. RSA2320 (SRS4604247) | 350 | 8463 | 520.3 | 9.9 | |
| Coemansia sp. RSA2320 (SRS4604247) | 433 | 7162 | 2772.9 | 52.7 | |
| Coemansia sp. RSA2320 (SRS4604247) | 453 | 6785 | 5065.5 | 96.3 | |
| Coemansia sp. RSA2320 (SRS4604247) | 514 | 5996 | 2807.8 | 53.4 | |
| Coemansia sp. RSA2320 (SRS4604247) | 538 | 5775 | 4305.7 | 81.9 | |
| Coemansia sp. RSA2320 (SRS4604247) | 565 | 5436 | 577.4 | 11.0 | |
| Coemansia sp. RSA2320 (SRS4604247) | 692 | 4597 | 1160.7 | 22.1 | |
| Coemansia sp. RSA2320 (SRS4604247) | 810 | 3983 | 1919.4 | 36.5 | 35.5 |

Illumina data from the SRA database was assembled using SPAdes. In SPAdes assemblies, contigs are called Nodes, and they are numbered in descending order of length. Rows highlighted in gray show the absolute coverage and length of a contig (usually Node 1) in the genome assembly of each species that was used as a proxy for the average coverage of its nuclear genome. For Cunninghamella blakesleeana VKA1, whose assembly consisted of mixed contigs from C. blakesleeana and M. guilliermondii, we used Node93, which was the largest non-rDNA and non-mitochondrial contig that hit proteins from Cunninghamellaceae species in BLASTX searches, as the proxy for average coverage of the C. blakesleeana nuclear genome. Many small contigs in assemblies are not VLEs; only contigs containing identifiable VLE genes are listed here and drawn in Figures 2 and 4.

**Table S2.** Estimated numbers of DNA polymerases in genome assemblies of Zoopagomycota species available at NCBI.

| NCBI accession prefix | Number of DNAPs | Species/strain | Clade (in Reynolds et al 2023) |
|---|---|---|---|
| JANBUG | 44 | Coemansia sp. BCRC 34301 | Kickxellales2 |
| JANBTZ | 39 | Coemansia sp. IMI 209128 | Kickxellales2 |
| JANBTX | 32 | Coemansia spiralis CBS 109367 | Kickxellales2 |
| JANBVB | 27 | Coemansia aciculifera CBS 190363 | Kickxellales2 |
| JANBQN | 27 | Coemansia sp. S146 | Kickxellales2 |
| JANBRM | 23 | Coemansia sp. RSA 455 | Kickxellales2 |
| JZJC | 17 | Coemansia reversa NRRL 1564 | Coemansia |
| JANBVA | 17 | Coemansia aciculifera CBS 516.66 | Kickxellales2 |
| JANBVC | 15 | Coemansia aciculifera BCRC 34181 | Kickxellales2 |
| JANBPH | 14 | Coemansia sp. IMI 209127 | Kickxellales1 |
| JANBSV | 14 | Coemansia sp. RSA 2049 | Kickxellales1 |
| JANBOT | 14 | Coemansia sp. RSA 2399 | Kickxellales1 |
| JANJYS | 12 | Coemansia sp. RSA 1646 | Kickxellales1 |
| JASJQH | 11 | Basidiobolus ranarum AG-B5 | --- |
| JASVEC | 11 | Conidiobolus obscurus 7217 | --- |
| JANBRJ | 11 | Coemansia sp. RSA 487 | Kickxellales1 |
| JANBOQ | 10 | Coemansia sp. RSA 1843 | Kickxellales1 |
| JANBTF | 9 | Coemansia sp. RSA 1813 | Kickxellales1 |
| JANBQR | 9 | Coemansia sp. RSA 986 | Kickxellales1 |
| JANBTA | 8 | Coemansia sp. RSA 1933 | Kickxellales1 |
| JANBSR | 8 | Coemansia sp. RSA 2320 | Kickxellales2 |
| JANBRZ | 8 | Coemansia sp. RSA 2673 | Kickxellales2 |
| JANBQF | 8 | Coemansia thaxteri IMI 214461 | Kickxellales2 |
| JANBTV | 7 | Coemansia sp. RSA 1085 | Coemansia |
| JANBSQ | 7 | Coemansia sp. RSA 2322 | Kickxellales2 |
| JANJYR | 7 | Coemansia sp. RSA 25 | Kickxellales2 |
| JANBQE | 7 | Coemansia thaxteri IMI 214463 | Kickxellales2 |
| JANBUW | 6 | Coemansia brasiliensis NRRL 1566 | Coemansia |
| JANBOL | 6 | Coemansia sp. RSA 1086 | Coemansia |
| JANBTS | 6 | Coemansia sp. RSA 1250 | Coemansia |
| JANBTO | 6 | Coemansia sp. RSA 1290 | Coemansia |
| JANBOO | 6 | Coemansia sp. RSA 1821 | Coemansia |
| JANBSC | 6 | Coemansia sp. RSA 2611 | Coemansia |
| JANBPD | 6 | Coemansia sp. RSA 989 | Coemansia |
| JANBPE | 6 | Coemansia sp. RSA 990 | Coemansia |
| JANBUD | 6 | Coemansia sp. Benny D115 | Kickxella |
| JANBTJ | 6 | Coemansia sp. RSA 1722 | Kickxella |
| JANBRK | 6 | Coemansia sp. RSA 485 | Kickxella |
| JANBPB | 6 | Coemansia sp. RSA 486 | Kickxella |
| JANBTT | 6 | Coemansia sp. RSA 1200 | Kickxellales1 |
| JANBTN | 6 | Coemansia sp. RSA 1358 | Kickxellales1 |
| JANBTW | 6 | Coemansia spiralis NRRL 3115 | Kickxellales1 |
| JANBQD | 6 | Coemansia umbellata BCRC 34882 | Kickxellales1 |
| JANBUE | 6 | Coemansia sp. BCRC 34962 | Kickxellales2 |
| JANBTL | 6 | Coemansia sp. RSA 1694 | Kickxellales2 |
| JANBSU | 6 | Coemansia sp. RSA 2050 | Kickxellales2 |
| JANBSN | 6 | Coemansia sp. RSA 2424 | Kickxellales2 |
| JANBPF | 6 | Coemansia sp. S17 | Kickxellales2 |
| JANBQJ | 6 | Coemansia sp. S3946 | Kickxellales2 |
| JANBTE | 5 | Coemansia sp. RSA 1822 | Coemansia |
| JANBRE | 5 | Coemansia sp. RSA 530 | Coemansia |
| JAOPJG | 5 | Dispira simplex RSA 1455 | Dimargaritales |
| JANBPG | 5 | Kickxella alabastrina Benny 63K | Kickxella |
| JANBTH | 5 | Coemansia sp. RSA 1804 | Kickxellales1 |
| JANBTD | 5 | Coemansia sp. RSA 1836 | Kickxellales2 |
| JANBQI | 5 | Coemansia sp. S610 | Kickxellales2 |
| WBMQ | 4 | Capillidium heterosporum CGMCC 3.15857 | --- |
| JAWQFY | 4 | Capillidium rhysosporum ATCC 12588 | --- |
| QTSX | 4 | Entomophthora muscae Berkeley | --- |
| LSSM | 4 | Smittium culicis ID-206-W2 | --- |
| MBFS | 4 | Smittium megazygosporum SC-DP-2 | --- |
| QZWR | 4 | Zoophagus insidians Zi-All | --- |
| JAIXMU | 4 | Coemansia mojavensis RSA 71 | Coemansia |
| JANBUB | 4 | Coemansia sp. D1744 | Coemansia |
| JANBTU | 4 | Coemansia sp. RSA 1199 | Coemansia |
| JANBOM | 4 | Coemansia sp. RSA 1591 | Coemansia |
| JANBON | 4 | Coemansia sp. RSA 1752 | Coemansia |
| JANBTI | 4 | Coemansia sp. RSA 1797 | Coemansia |
| JANBTC | 4 | Coemansia sp. RSA 1853 | Coemansia |
| JANBOR | 4 | Coemansia sp. RSA 1938 | Coemansia |
| JANBSW | 4 | Coemansia sp. RSA 1972 | Coemansia |
| JANBSM | 4 | Coemansia sp. RSA 2440 | Coemansia |
| JANBSB | 4 | Coemansia sp. RSA 2618 | Coemansia |
| JANBRT | 4 | Coemansia sp. RSA 353 | Coemansia |
| JANBRR | 4 | Coemansia sp. RSA 370 | Coemansia |
| JANBRQ | 4 | Coemansia sp. RSA 371 | Coemansia |
| JANBRL | 4 | Coemansia sp. RSA 475 | Coemansia |
| JANBRI | 4 | Coemansia sp. RSA 518 | Coemansia |
| JANBRG | 4 | Coemansia sp. RSA 521 | Coemansia |
| JANBRF | 4 | Coemansia sp. RSA 522 | Coemansia |
| JANBRD | 4 | Coemansia sp. RSA 532 | Coemansia |
| JANBQY | 4 | Coemansia sp. RSA 564 | Coemansia |
| JANBPC | 4 | Coemansia sp. RSA 638 | Coemansia |
| JANBQV | 4 | Coemansia sp. RSA 720 | Coemansia |
| JANBQU | 4 | Coemansia sp. RSA 788 | Coemansia |
| JANBTY | 4 | Coemansia spiralis CBS 108938 | Coemansia |
| JANBPT | 4 | Tieghemiomyces parasiticus RSA 861 | Dimargaritales |
| JANBOH | 4 | Coemansia asiatica NBRC 105413 | Kickxella |
| JANBUS | 4 | Coemansia erecta IMI 279145 | Kickxella |
| JANBTK | 4 | Coemansia sp. RSA 1721 | Kickxella |
| JANBPX | 4 | Kickxella alabastrina RSA 2140 | Kickxella |
| JANBSX | 4 | Coemansia sp. RSA 1939 | Kickxellales1 |
| JANBUZ | 4 | Coemansia aciculifera NRRL 2694 | Kickxellales2 |
| JANBUY | 4 | Coemansia aciculifera RSA 476 | Kickxellales2 |
| JANBUP | 4 | Coemansia furcata CBS 102833 | Kickxellales2 |
| JANBUH | 4 | Coemansia pectinata BCRC 34297 | Kickxellales2 |
| JANBST | 4 | Coemansia sp. RSA 2052 | Kickxellales2 |
| JANBSO | 4 | Coemansia sp. RSA 2337 | Kickxellales2 |
| JANBSI | 4 | Coemansia sp. RSA 2530 | Kickxellales2 |
| JANBSH | 4 | Coemansia sp. RSA 2531 | Kickxellales2 |
| JANBOX | 4 | Coemansia sp. RSA 2675 | Kickxellales2 |
| JANBRY | 4 | Coemansia sp. RSA 2681 | Kickxellales2 |
| JANBQS | 4 | Coemansia sp. RSA 922 | Kickxellales2 |
| JANBQP | 4 | Coemansia sp. S100 | Kickxellales2 |

| | | | |
|---|---|---|---|
| JANBQO | 4 | Coemansia sp. S142-1 | Kickxellales2 |
| JANBQL | 4 | Coemansia sp. S16 | Kickxellales2 |
| JANBQH | 4 | Coemansia sp. S680 | Kickxellales2 |
| JANBPZ | 4 | Dipsacomyces acuminosporus NRRL 2925 | Kickxellalesout |
| JANBPV | 4 | Linderina pennispora NRRL 3781 | Kickxellalesout |
| JBNZCG | 3 | Conidiobolus coronatus 20220622F | --- |
| MCFD | 3 | Linderina pennispora ATCC 12442 DL89 | --- |
| LSSN | 3 | Smittium culicis GSMNP | --- |
| JAIXMF | 3 | Syncephalis plumigaleata NRRL S24 | --- |
| LSSK | 3 | Zancudomyces culisetae COL-18-3 | --- |
| JANBOI | 3 | Coemansia biformis BCRC 34381 | Coemansia |
| JANBUT | 3 | Coemansia erecta CBS 939.97 | Coemansia |
| JANBOJ | 3 | Coemansia erecta NBRC 32514 | Coemansia |
| JANBUN | 3 | Coemansia helicoidea BCRC 34780 | Coemansia |
| JANBUC | 3 | Coemansia sp. Cherry 401B | Coemansia |
| JANBTP | 3 | Coemansia sp. RSA 1287 | Coemansia |
| JANBTM | 3 | Coemansia sp. RSA 1365 | Coemansia |
| JANBTG | 3 | Coemansia sp. RSA 1807 | Coemansia |
| JANBTB | 3 | Coemansia sp. RSA 1878 | Coemansia |
| JANBSS | 3 | Coemansia sp. RSA 2131 | Coemansia |
| JANBOS | 3 | Coemansia sp. RSA 2167 | Coemansia |
| JANBSP | 3 | Coemansia sp. RSA 2336 | Coemansia |
| JANBSL | 3 | Coemansia sp. RSA 2522 | Coemansia |
| JANBRX | 3 | Coemansia sp. RSA 2702 | Coemansia |
| JANBRW | 3 | Coemansia sp. RSA 2704 | Coemansia |
| JANBRV | 3 | Coemansia sp. RSA 2705 | Coemansia |
| JANBRU | 3 | Coemansia sp. RSA 2706 | Coemansia |
| JANBRS | 3 | Coemansia sp. RSA 355 | Coemansia |
| JANBRO | 3 | Coemansia sp. RSA 451 | Coemansia |
| JANBRH | 3 | Coemansia sp. RSA 520 | Coemansia |
| JANBRB | 3 | Coemansia sp. RSA 552 | Coemansia |
| JANBRA | 3 | Coemansia sp. RSA 560 | Coemansia |
| JANBQZ | 3 | Coemansia sp. RSA 562 | Coemansia |
| JANBQX | 3 | Coemansia sp. RSA 637 | Coemansia |
| JANBQT | 3 | Coemansia sp. RSA 921 | Coemansia |
| JANBQQ | 3 | Coemansia sp. RSA 988 | Coemansia |
| JAMZIH | 3 | Spiromyces aspiralis RSA 2271 | Harpellales |
| JANBUX | 3 | Coemansia asiatica BCRC 34628 | Kickxella |
| JANBUU | 3 | Coemansia erecta CBS 336.87 | Kickxella |
| JANBUA | 3 | Coemansia sp. IMI 203386 | Kickxella |
| JANBTQ | 3 | Coemansia sp. RSA 1286 | Kickxella |
| JANBOV | 3 | Coemansia sp. RSA 2598 | Kickxella |
| JANBOW | 3 | Coemansia sp. RSA 2599 | Kickxella |
| JANBTR | 3 | Coemansia sp. RSA 1285 | Kickxellales1 |
| JANBSG | 3 | Coemansia sp. RSA 2559 | Kickxellales1 |
| JAIXMV | 3 | Coemansia spiralis RSA 1278 | Kickxellales1 |
| JANBUQ | 3 | Coemansia furcata BCRC 34190 | Kickxellales2 |
| JANBUK | 3 | Coemansia linderi BCRC 34191 | Kickxellales2 |
| JANBUR | 3 | Coemansia sp. 'formosensis' NRRL 5531 | Kickxellales2 |
| JANBSA | 3 | Coemansia sp. RSA 2671 | Kickxellales2 |
| JANBQM | 3 | Coemansia sp. S155-1 | Kickxellales2 |
| JANBQK | 3 | Coemansia sp. S2 | Kickxellales2 |
| JANBPW | 3 | Linderina macrospora NRRL 5244 | Kickxellalesout |
| JAIXMM | 3 | Martensiomyces pterosporus CBS 209.56 | Kickxellalesout |
| QZWU | 2 | Acaulopage tetraceros T-All | --- |
| MCFE | 2 | Basidiobolus meristosporus CBS 931.73 | --- |
| QRFA | 2 | Dimargaris cristalligena RSA 468 | --- |
| MBFT | 2 | Furculomyces boomerangus AUS-77-4 | --- |
| JAKSZP | 2 | Massospora cicadina MCPNR19 | --- |
| QKRY | 2 | Massospora platypediae k141 | --- |
| JAHVYE | 2 | Neoconidiobolus thromboides FSU 785 K502 | --- |
| MBFU | 2 | Smittium angustum AUS-126-30 | --- |
| JBJCJL | 2 | Smittium minutisporum WKRb | --- |
| LSSL | 2 | Smittium mucronatum ALG-7-W6 | --- |
| MBFR | 2 | Smittium simulii SWE-8-4 | --- |
| SSNV | 2 | Stylopage hadra SOG | --- |
| JANBUL | 2 | Coemansia javaensis NBRC 105414 | Coemansia |
| JANBUJ | 2 | Coemansia nantahalensis CBS 109366 | Coemansia |
| JANBUI | 2 | Coemansia nantahalensis CBS 109400 | Coemansia |
| JANBOP | 2 | Coemansia sp. RSA 1824 | Coemansia |
| JANBSZ | 2 | Coemansia sp. RSA 1935 | Coemansia |
| JANBSY | 2 | Coemansia sp. RSA 1937 | Coemansia |
| JANBOU | 2 | Coemansia sp. RSA 2523 | Coemansia |
| JANBSK | 2 | Coemansia sp. RSA 2524 | Coemansia |
| JANBSJ | 2 | Coemansia sp. RSA 2526 | Coemansia |
| JANBSD | 2 | Coemansia sp. RSA 2610 | Coemansia |
| JANBPA | 2 | Coemansia sp. RSA 2711 | Coemansia |
| JANBRN | 2 | Coemansia sp. RSA 454 | Coemansia |
| JANBRC | 2 | Coemansia sp. RSA 551 | Coemansia |
| JANBQW | 2 | Coemansia sp. RSA 678 | Coemansia |
| JANBQC | 2 | Dimargaris cristalligena RSA 1219 | Dimargaritales |
| JANBQB | 2 | Dimargaris verticillata RSA 567 | Dimargaritales |
| JANBQA | 2 | Dimargaris xerosporica NRRL 3178 | Dimargaritales |
| JANBPY | 2 | Dispira parvispora RSA 1196 | Dimargaritales |
| JAJHGN | 2 | Ramicandelaber brevisporus CBS 109374 | Dimargaritales |
| JANBPU | 2 | Mycoemilia scoparia NBRC 100468 | Harpellales |
| JANBUM | 2 | Coemansia interrupta BCRC 34489 | Kickxella |
| JANBSF | 2 | Coemansia sp. RSA 2603 | Kickxella |
| JANBSE | 2 | Coemansia sp. RSA 2607 | Kickxella |
| JANBOY | 2 | Coemansia sp. RSA 2703 | Kickxella |
| JALLKQ | 2 | Kickxella alabastrina RSA 675 | Kickxella |
| JANBUV | 2 | Coemansia erecta BCRC 34228 | Kickxellales1 |
| QZWT | 1 | Cochlonema odontosperma E-All | --- |
| QMCF | 1 | Massospora cicadina MICH 231384 | --- |
| QPFT | 1 | Piptocephalis cylindrospora RSA 2659 | --- |
| JAMKHV | 1 | Piptocephalis tieghemiana RSA 1565 | --- |
| QUVU | 1 | Thamnocephalis sphaerospora RSA 1356 | --- |
| JANBUO | 1 | Coemansia guatemalensis NRRL 1565 | Coemansia |
| JANBOZ | 1 | Coemansia sp. RSA 2708 | Coemansia |
| JANBUF | 1 | Coemansia sp. BCRC 34490 | Kickxellales1 |
| JANBOK | 1 | Coemansia sp. Benny D160-2 | Kickxellales1 |
| JANBRP | 1 | Coemansia sp. RSA 376 | Kickxellales2 |
| JANBQG | 1 | Coemansia sp. S85 | Kickxellales2 |
